# Expanding the Genetic Code of an Animal with Two Non-Canonical Amino Acids

**DOI:** 10.64898/2026.01.20.700547

**Authors:** Jose Javier Vazquez Rodriguez, Sebastian Greiss

## Abstract

Expanding the genetic code of multicellular organisms for the incorporation of non-canonical amino acids has thus far been limited to a single non-canonical amino acid at a time. To overcome this limitation, we establish the pyrrolysyl-tRNA-synthetase (PylRS)/tRNA^Pyl^ pair from *Methanomethylophilus alvus* in *Caenorhabditis elegans* and show that it is highly efficient and mutually orthogonal to the *Methanosarcina mazei* PylRS/tRNA^Pyl^ pair previously established in the worm. We furthermore establish a set of mutually orthogonal blank codons for use in *C. elegans* that allow the simultaneous and independent incorporation of two ncAAs for the first time in an animal. We then engineer the non-canonical amino acid specificity of *M. alvus* PylRS to incorporate several photocaged ncAAs. Taken together, this allows expression of photocaged variants of FLP and Cre using two different photocaged amino acids, enabling us to optically control the expression of two target genes.

## INTRODUCTION

Genetic code expansion (GCE) allows the co-translational and site-specific incorporation of non-canonical amino acids (ncAA) into proteins, augmenting the repertoire of amino acid building blocks available to living systems. A wide range of diverse ncAAs with functionalities such as bioorthogonal handles, post translational modifications, crosslinkers, spectroscopic probes, and photocages have been introduced into proteins in a variety of systems including bacteria, eukaryotic cells and multicellular organisms^1–8^.

GCE is achieved through the use of orthogonal aminoacyl-tRNA-synthetase (aaRS)/tRNA pairs. The aaRS specifically aminoacylates its cognate tRNA with an ncAA that is then co-translationally incorporated at a blank codon not assigned to a canonical amino acid. Pairs that have been used for GCE include the *Methanocaldococcus jannaschii* tyrosyl-tRNA synthetase (*Mj*TyrRS)/tRNA^Tyr^ pair, which is orthogonal in bacteria, the *Ec*TyrRS/tRNA^Tyr^ and *Ec*LeuRS/tRNA^Leu^ pairs from *E. coli*, which are orthogonal in eukaryotic cells, and pyrrolysyl-tRNA-synthetase (PylRS)/tRNA^Pyl^ pairs, which are orthogonal in both *E. coli* and in eukaryotic cells.

Of the available aaRS/tRNA pairs, those based on PylRS/tRNA^Pyl^ are currently the most widely used, owing to their efficiency and versatility. PylRS/tRNA^Pyl^ pairs are found in a variety of methanogenic bacteria and archaea, where they serve to direct the incorporation of pyrrolysine, the 22nd naturally occurring proteinogenic amino acid. Wild type PylRS can however be engineered through directed evolution, usually performed in *E. coli*, to alter its substrate specificity, and evolved variants can be directly applied in eukaryotes. The first PylRS family members used for GCE, taken from methanogenic *Methanosarcina* species, have to date been evolved to recognise a wide array of diverse ncAAs^4^. Additionally, unlike most aaRSs, PylRSs do not use the tRNA^Pyl^ anticodon as an identity element^9^, making it possible to change codon specificity without the need to engineer the PylRS. The flexibility of the PylRS/tRNA^Pyl^ system has allowed the use of stop codons other than the most commonly used TAG codon, as well as quadruplet codons^10–12^, sense codons^13^, and codons that include non-canonical bases^14^.

Strategies have been explored to enable the incorporation of multiple ncAAs independently in the same cell. This requires aaRS/tRNA pairs that are orthogonal to the endogenous pairs and also orthogonal to each other. One aaRS/tRNA pair and one codon are required for each ncAA to be incorporated. Examples include the use of a PylRS/tRNA^Pyl^ pair together with the *Mj*TyrRS/tRNA^Tyr^ pair to incorporate two ncAAs in *E. coli*^15^, and the use of a PylRS/tRNA^Pyl^ pair together with the *Ec*TyrRS/tRNA^Tyr^ pair in mammalian cell culture^16^. However, unlike PylRS/tRNA^Pyl^ pairs, these additional aaRS/tRNA pairs used are limited in the codons that they can decode without extensive aaRS engineering^4^, and are not orthogonal in both bacteria and eukaryotes, limiting the application of any system only to the host in which it is developed.

More recently, these limitations have been overcome with the discovery of distinct PylRS subclasses that can be used to establish mutually orthogonal PylRS/tRNA^Pyl^ pairs^17–21^. The PylRSs from *Methanosarcina mazei* (*Mm*) and *Methanosarcina barkeri* (*Mb*), the most commonly used for GCE, are archetypal of the PylRS N-C class, which consists of enzymes that contain a catalytic C-terminal domain and an essential N-terminal domain that binds the T-arm and variable loop of the tRNA. A second class, called ΔNPylRS, consists of proteins that contain only the catalytic C-terminal domain and lack the N-terminal domain. The PylRSs from *Methanomethylophilus alvus* (*Ma*) and *Methanogenic archaeon ISO4-G1* (*G1*) are archetypal of this class. Initial experiments exploring orthogonality between members of the different subclasses showed that, while *Ma*PylRS and *G1*PylRS do not interact with *Mm*tRNA^Pyl^, their respective tRNA^Pyl^s do interact with *Mm*PylRS^17,20,21^.It was however found that mutations and insertions could be introduced into the variable loop of *Ma* and *G1* tRNA^Pyl^ to disrupt their interaction with *Mm*PylRS, while preserving the interaction with their respective PylRS. This engineered mutual orthogonality between PylRS/tRNA^Pyl^ pairs has allowed the use of the highly efficient PylRS/tRNA^Pyl^ system for the incorporation of more than one ncAA^17,20–22^.

Mutually orthogonal PylRS/tRNA^Pyl^ pairs have been used to incorporate two ncAAs in eukaryotic cells, and up to 3 ncAAs in bacteria^17,18,20–22^. However, thus far, the genetic code of a multicellular organism has not been expanded beyond a single ncAA at a time.

Here, we report the first instance of double ncAA incorporation in an animal, the nematode worm *C. elegans*. We first establish the *Ma*PylRS/tRNA^Pyl^ and *G1*PylRS/tRNA^Pyl^ pairs in *C. elegans* and determine their orthogonality to *Mm*PylRS/tRNA^Pyl^ variants. We then introduce new blank triplet and quadruplet codons in *C. elegans*, allowing us to independently incorporate two ncAAs. We go on to engineer the newly established *Ma*PylRS pair to recognise several different photocaged amino acids. Finally, we apply our new double incorporation system to concurrently express photocaged versions of FLP and Cre DNA recombinases using different ncAAs. This allows us to optically control expression of two target genes using different wavelengths.

## RESULTS

### Establishing *M. alvus* and *ISO4-G1* PylRS/tRNA^Pyl^ pairs in *C. elegans*

The *Ma*PylRS/tRNA^Pyl^ and *G1*PylRS/tRNA^Pyl^ pairs, engineered to be orthogonal with respect to the *Mm*PylRS/tRNA^Pyl^ pair, are highly active and orthogonal in *E. coli* and mammalian cell culture^17,20,21^. Given this, we sought to assess their activity and orthogonality in *C. elegans*. Specifically, we chose the *Ma*PylRS/tRNA^Ma6^, *Ma*PylRS/tRNA^Ma10^, and *G1*PylRS/tRNA^G1hyb^ pairs, as these were shown to be the most efficient of the orthogonalised pairs^17,20,21^.

We generated transgenic *C. elegans* strains by co-transforming constructs encoding each PylRS, its respective tRNA^Pyl^, and a fluorescent GFP:: mCherry reporter bearing an amber (TAG) stop codon within a linker between the GFP and mCherry genes^23^ (**Fig. 1a**). For this reporter, ncAA incorporation in response to the TAG codon leads to translation of the full-length GFP::mCherry fusion protein, while translation in the absence of ncAA, or unsuccessful incorporation leads to termination at the TAG stop codon and expression of GFP alone. Relative efficiency of ncAA incorporation can then be determined by the ratio of red mCherry fluorescence to green GFP fluorescence. Additionally, the reporter contains a C-terminal nuclear localisation sequence to concentrate the full-length product in the nucleus, as well as an HA tag for easy detection by western blot.

**Figure 1.**
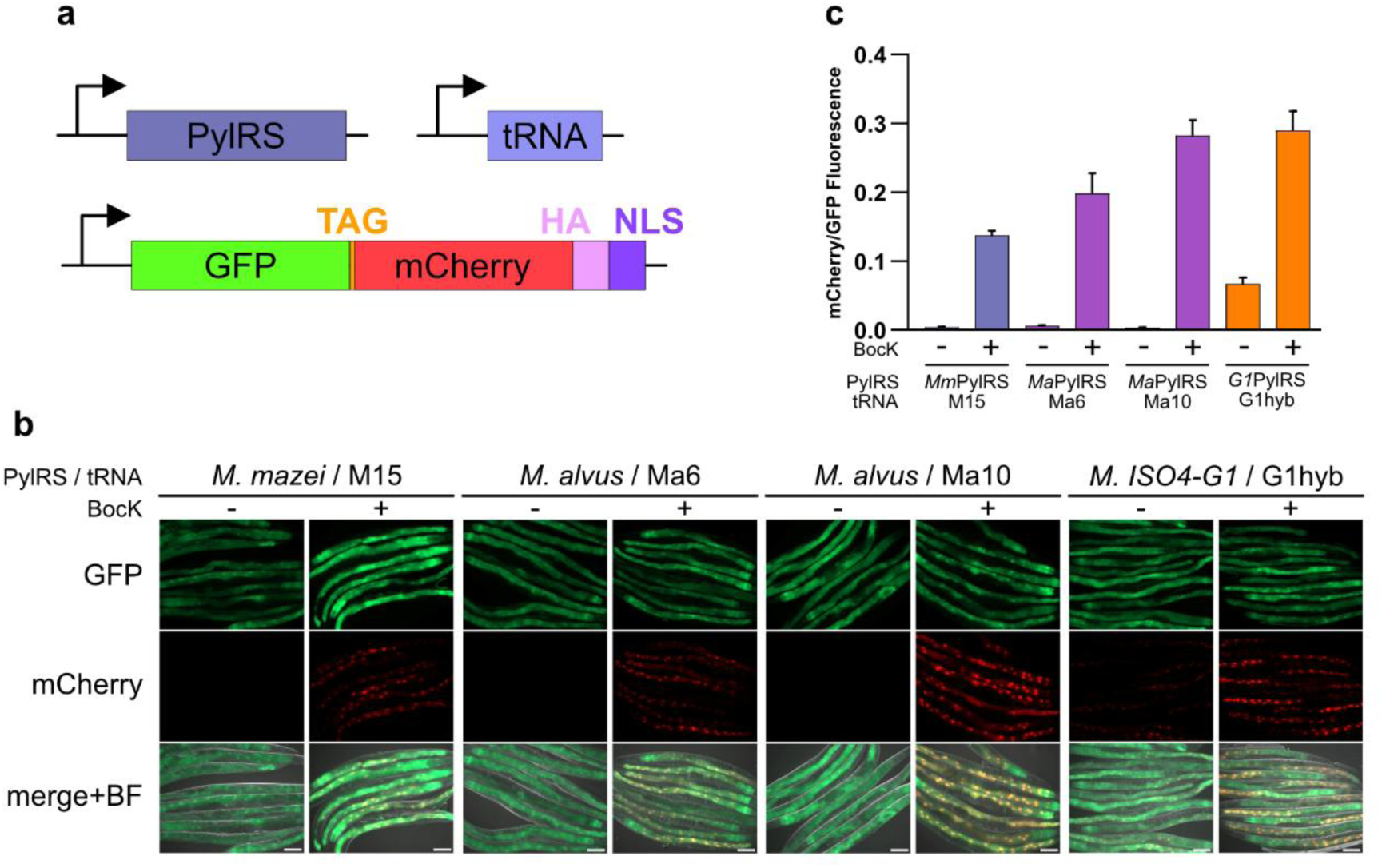
Additional PylRS/tRNA^Pyl^ pairs. **a** Constructs transformed into *C. elegans* for assaying the activity of PylRS/tRNA^Pyl^ variants. Reporter consists of a GFP gene separated from an mCherry gene by an amber stop codon. An HA tag and nuclear localisation sequence follow downstream of the mCherry. **b** Fluorescence microscopy images of worms expressing select PylRS/tRNA^Pyl^ variants and grown in the presence or absence of BocK. BF stands for brightfield. Scale bars represent 100 μm. **c** mCherry to GFP fluorescence ratio of worms expressing select PylRS/tRNA^Pyl^ variants and grown in the presence or absence of BocK. Error bars represent the SEM.

To assess ncAA incorporation, we grew transgenic animals from the L4 larval stage for 24 hours on NGM agar plates supplemented with 1mM boc-lysine (BocK), a known substrate for PylRSs^24^, or in the absence of ncAA. We found that all PylRS/tRNA^Pyl^ pairs tested showed clearly visible nuclear mCherry fluorescence, indicative of efficient BocK incorporation. Neither the *Ma*PylRS/tRNA^Ma6^ or *Ma*PylRS/tRNA^Ma10^ pairs showed any mCherry fluorescence in the absence of the ncAA. In contrast, the *G1*PylRS/tRNA^G1hyb^ pair displayed substantial mCherry fluorescence even in the absence of ncAA, suggesting incomplete orthogonality in *C. elegans* (**Fig. 1b**).

We then quantified the relative incorporation efficiency of each pair and compared it to the previously established *Mm*PylRS/tRNA^M15^ system^25,26^, which consists of the tRNA^M15^ variant optimised for activity in eukaryotic systems^16^, and an *Mm*PylRS with an added N-terminal nuclear export signal that promotes cytoplasmic localisation^26,27^. We found that the *Ma*PylRS/tRNA^Ma6^ and *Ma*PylRS/tRNA^Ma10^ pairs were both highly active, showing relative mean incorporation levels of 145% and 205%, respectively, as compared to the established *Mm*PylRS/tRNA^M15^ pair (**Fig. 1c**). *G1*PylRS/tRNA^G1hyb^ was also highly active, showing 211% of *Mm*PylRS/tRNA^M15^ in the presence of ncAA, albeit with the limitation that animals also showed clearly detectable red fluorescence even in the absence of ncAA.

### Mutual orthogonality of PylRS/tRNA^Pyl^ pairs

Since incorporation of two ncAAs requires two mutually orthogonal aaRS/tRNA pairs, we next determined cross reactivity between PylRSs and tRNA^Pyl^s from the different species. For this, we generated transgenic strains expressing all possible PylRS/tRNA^Pyl^ combinations along with the GFP:: mCherry reporter (**Fig. 2a**). Here we also included tRNA^C15^, an additional optimised tRNA variant distinct from tRNA^M15^, which we previously found to also be highly active in combination with *Mm*PylRS in *C. elegans*^16,26^.

**Figure 2.**
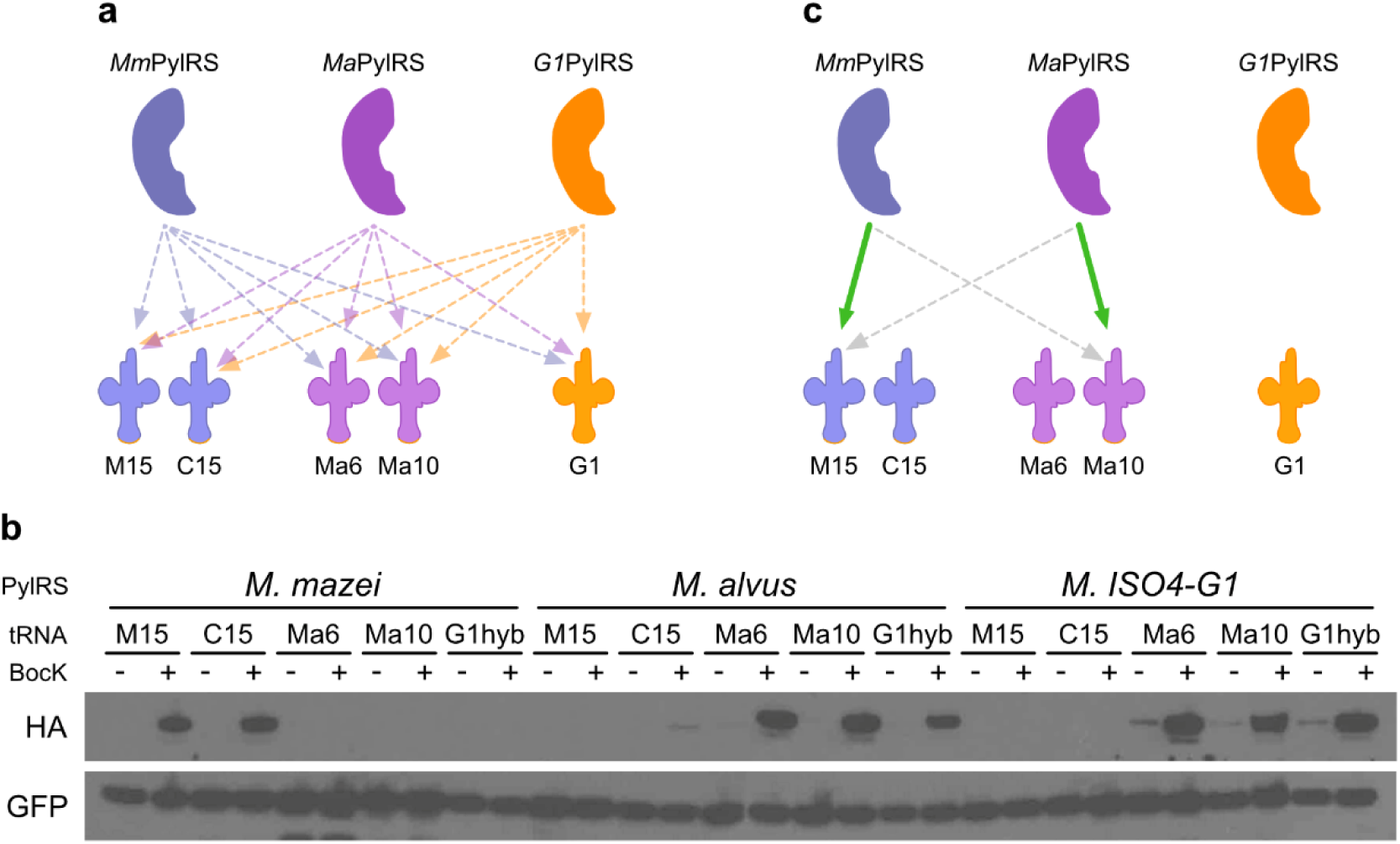
Mutually orthogonal PylRS/tRNA^Pyl^ pairs. **a** Schematic of all the possible PylRS and tRNA^Pyl^ combinations that were tested to identify two mutually orthogonal pairs. **b** Western blot against the HA tag present only in the full-length reporter GFP::mCherry used to identify interacting pairs. GFP was used as a loading control. **c** Mutually orthogonal PylRS/tRNA^Pyl^ pairs chosen for further experiments. Green arrows represent interaction, broken grey arrows represent no interaction.

When we grew transgenic animals on NGM agar plates supplemented with ncAA , we found strong expression of full-length GFP::mCherry reporter protein, indicative of non-orthogonality, in strains expressing *Ma*PylRS/tRNA^G1hyb^ as well as in the reverse combinations of *G1*PylRS/tRNA^Ma6^ and *G1*PylRS/tRNA^Ma10^ (**Fig. 2b**). We furthermore saw a weak signal in the combination *Ma*PylRS/tRNA^C15^ suggesting incomplete orthogonality. In contrast, we could detect no ncAA incorporation in strains expressing the combinations *Ma*PylRS/tRNA^M15^, *Mm*PylRS/tRNA^Ma6^ and *Mm*PylRS/tRNA^Ma10^. While we could also not detect any ncAA incorporation in the strain expressing the *Mm*PylRS/tRNA^G1hyb^ pairing, we did find clearly detectable GFP::mCherry reporter produced in the *G1*PylRS/tRNA^G1hyb^ strain in the absence of ncAA (**Fig. 1b, c**; **Fig, 2b**), and therefore eliminated this pair from further use.

We concluded that *Mm*PylRS/tRNA^M15^ in combination with either *Ma*PylRS/tRNA^Ma6^ or *Ma*PylRS/tRNA^Ma10^ could serve as two mutually orthogonal pairs in *C. elegans*. Given that the *Ma*PylRS/tRNA^Ma10^ pair was more efficient than the *Ma*PylRS/tRNA^Ma6^ pair (**Fig. 1c**), only it was used in further experiments (**Fig. 2c**).

### Establishing mutually orthogonal codons

Two codons are required to direct the incorporation of two ncAAs. These codons should be blank, i.e. not assigned to any other amino acid, and mutually orthogonal. In *C. elegans*, the only codons established thus far are the amber stop codon TAG and the quadruplet codon TAGA. Since TAG and TAGA are, however, not fully mutually orthogonal^25^, we set out to establish additional orthogonal codons for use in *C. elegans*.

In bacteria and mammalian cell culture, double incorporation has been achieved using two stop codons, or combinations of stop codons with quadruplet codons^11,15–17^. We therefore explored incorporation efficiency and orthogonality in *C. elegans* of the three stop codons, as well as several quadruplet codons.

To explore stop codons as a source of additional mutually orthogonal blank codons, we decided to first determine whether the stop codons could be orthogonally decoded, meaning that a tRNA^Pyl^ bearing the anticodon complementary to a stop codon does not decode either of the other two stop codons, as reports of their mutual orthogonality in the literature are conflicting^28^. Using the *Mm*PylRS/tRNA^M15^ pair, we generated transgenic strains, each expressing one of the nine possible tRNA anticodon to reporter codon combinations: tRNA^M15^ variants bearing either the CUA, UUA, or UCA anticodon, and the GFP::mCherry reporter bearing either TAG, TAA, or TGA between the GFP and mCherry genes. When we grew these animals in the presence of ncAA we found that, in all cases, the full-length GFP::mCherry reporter was only produced in the strains expressing the complementary codon-anticodon pairings, confirming that the three stop codons can be decoded and that they are mutually orthogonal in *C. elegans* (**Fig. 3a, b**).

**Figure 3.**
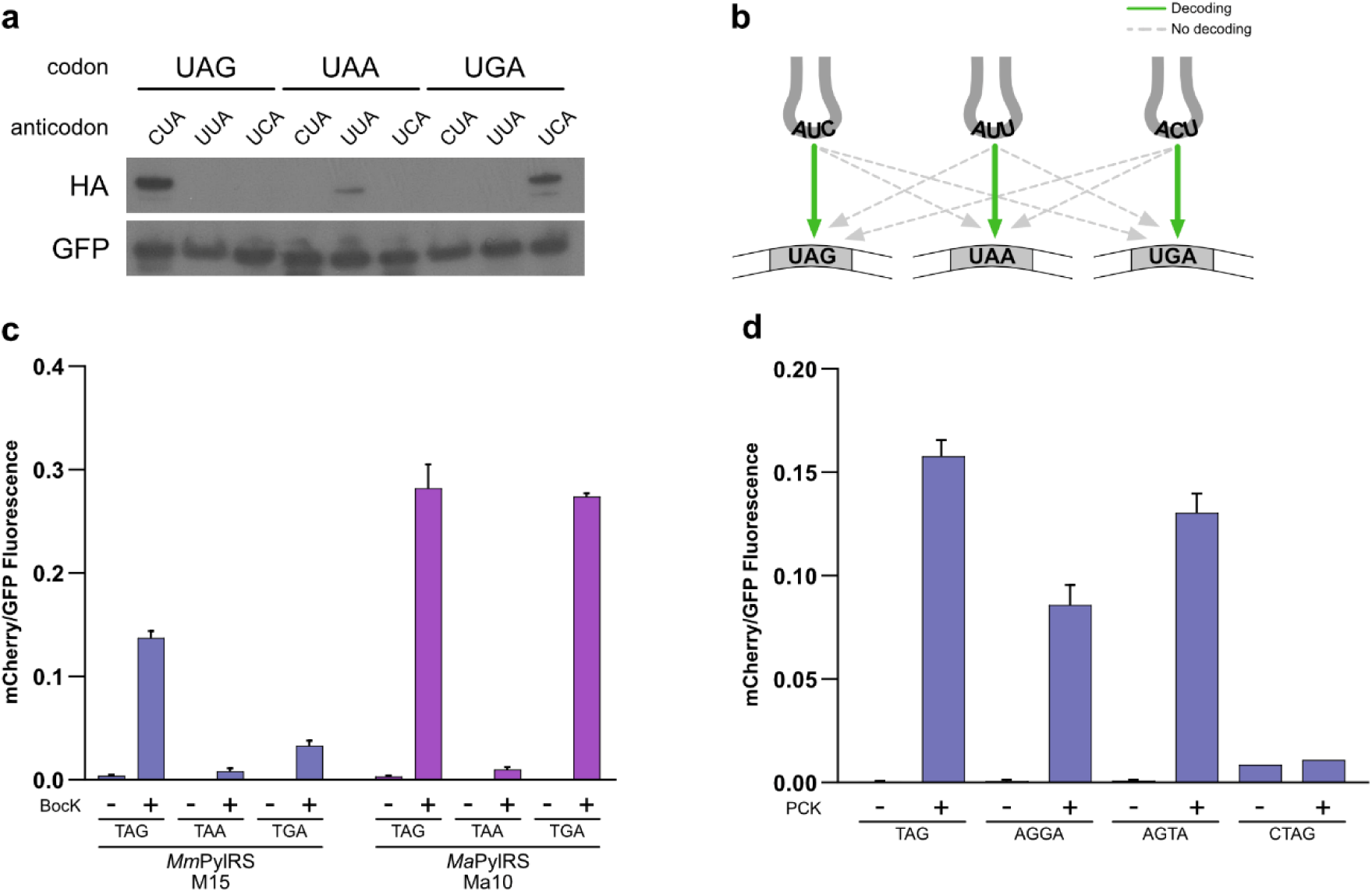
Mutually orthogonal codons. **a** Western blot against the HA tag present in the full-length GFP::mCherry reporter used to identify interacting codon anticodon partners. GFP was used as a loading control. **b** Schematic summary of results from **a**. Green arrows represent interaction, broken grey arrows represent no interaction. **c** mCherry to GFP fluorescence ratio of worms expressing a PylRS/tRNA^Pyl^ pair to suppress the respective stop codon in the GFP::mCherry reporter. Error bars represent the SEM. **d** mCherry to GFP fluorescence ratio of worms decoding the TAG, AGGA, AGTA, or CTAG codon in between the GFP and mCherry gene. Error bars represent the SEM.

Since all three stop codons could therefore potentially serve to incorporate two ncAAs, we proceeded to determine their efficiency when decoded by tRNA^M15^ or tRNA^Ma10^. We generated transgenic strains to incorporate ncAA into the GFP::mCherry reporter in response to each of the three stop codons using either the *Ma*PylRS/tRNA^Ma10^ or the *Mm*PylRS/tRNA^M15^ pairs. For both pairs we found that ncAA incorporation was possible for all three stop codons, and that TAA was the least efficient codon. For the *M. mazei* pair, incorporation was most efficient at the TAG codon, being 4.2-fold higher than at TGA. In contrast, the *M. alvus* pair showed high incorporation efficiency at both TAG and TGA codons, in both cases outperforming the *M. mazei* tRNA^M15^ (**Fig. 3c**).

For *M. mazei* tRNA^Pyl^, several different optimised quadruplet-decoding variants have been developed by modifying the anticodon loop through directed evolution. We have previously shown that such evolved anticodon loops can be combined with efficiency-enhancing mutations in the tRNA scaffold to create optimised hybrid tRNA variants ^25^. We therefore constructed hybrid tRNA variants consisting of the tRNA^M15^ scaffold fused to anticodon loops evolved for decoding AGGA^10^, AGTA^11^, CTAG^11^ quadruplet codons.

We generated transgenic strains expressing each quadruplet-decoding tRNA^M15^ variant together with a GFP::mCherry reporter bearing the complementary quadruplet codon inserted between the GFP and mCherry genes. When grown on NGM agar plates supplemented with 1 mM ncAA, we found clear mCherry fluorescence for AGGA and AGTA, with efficiencies of 54.4% and 82.7% as compared to the level reached when using a TAG codon for incorporation (**Fig. 3d**). We found no significant mCherry signal above background for the CTAG reporter. These results placed the AGGA and AGTA codons as potential additional blank codons.

Taken together, these experiments identified the TAG or TGA codons decoded by the *Ma*PylRS/tRNA^Ma10^ pair and the codons TAG, AGGA, or AGTA decoded by the *Mm*PylRS/tRNA^M15^ pair as suitable for double incorporation. We settled on the combination of the two pairs showing the highest incorporation efficiencies for all further double incorporation experiments: TAG decoded by *Mm*PylRS/tRNA^M15^ and TGA decoded by *Ma*PylRS/tRNA^Ma10^.

### Independent incorporation of two ncAAs

Having established mutually orthogonal PylRS/tRNA^Pyl^ pairs and mutually orthogonal codons to direct ncAA incorporation, we proceeded to generate transgenic *C. elegans* strains combining the genetic elements required for incorporation of two ncAAs. We selected the *Ma*PylRS/tRNA^Ma10^ pair to incorporate BocK, and the *Mm*PCKRS/tRNA^M15^ pair to incorporate photocaged lysine (PCK). *Mm*PCKRS is an *Mm*PylRS variant developed to incorporate PCK^29^, and is highly efficient in *C. elegans*^26^.

*Ma*PylRS and *Mm*PCKRS were co-expressed from an artificial *C. elegans* operon driven by the ubiquitous *sur-5p* promoter, and the tRNAs were expressed from separate expression cassettes each driven by a copy of the *rpr-1p* promoter. To assay ncAA incorporation, we again used a fluorescent GFP::mCherry reporter, albeit modified to contain a second blank codon. In addition to the TAG in the linker region between the GFP and mCherry genes to direct incorporation of PCK by *Mm*PCKRS/tRNA^M15^ , we inserted a TGA codon replacing the codon at position D212 in mCherry^30^ to direct incorporation of BocK by *Ma*PylRS/tRNA^Ma10^ (**Fig. 4a**).

**Figure 4.**
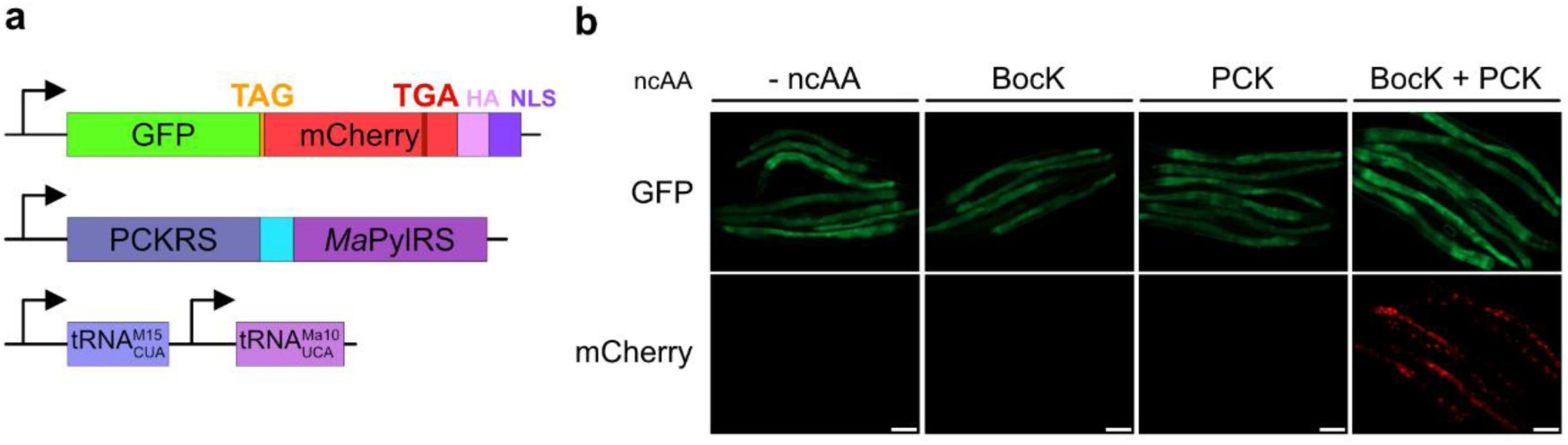
Double incorporation of the ncAAs PCK and BocK. **a** Schematic of genetic constructs used to create *C. elegans* strains. The light blue rectangle between *Mm*PCKRS and *Ma*PylRS represents the intergenic region between *C. elegans gpd-2* and *gpd-3* used to form a synthetic operon. **b** Fluorescence microscopy images of worms expressing the two mutually orthogonal PylRS/tRNA^Pyl^ pairs to decode TAG and TGA following growth in the presence and absence of the ncAAs. Scale bars represent 100 μm.

Transgenic animals were grown from the L4 larval stage for 24 hours on NGM agar plates supplemented either with 1 mM PCK alone, 1 mM BocK alone, 1 mM PCK and BocK, or no ncAA. We detected no mCherry fluorescence in animals grown in the absence of ncAA or in the presence of only a single ncAA. In contrast, animals grown in the presence of both ncAAs displayed clearly visible mCherry fluorescence, indicative of production of the full-length GFP::mCherry reporter resulting from ncAA incorporation at both TAG and TGA (**Fig. 4b**). This demonstrates the incorporation of two ncAAs into a single protein in an animal using mutually orthogonal PylRS/tRNA^Pyl^ pairs and mutually orthogonal codons.

### Engineering *M. alvus* PylRS to expand its ncAA repertoire

In nature, the substrate of wild type PylRS is pyrrolysine, and although PylRSs are to some extent promiscuous^31^, they cannot incorporate the majority of the ncAA repertoire currently available for GCE. To expand their utility, they must therefore be engineered to modify their substrate specificity and allow the incorporation of the wide array of available ncAAs. This is most often achieved by directed evolution of the active site, which requires the generation of randomised variant libraries followed by laborious rounds of positive and negative selection^32^. By this approach, many variants of *Mm*PylRS and *Mb*PylRS have been engineered to incorporate a large variety of ncAAs^4^. Since *Mm*PylRS and *Mb*PylRS are closely related, active site mutations engineered for one enzyme can normally be transferred to the other to grant it specificity for a given ncAA^24^. The same does however often not hold true for transplanting active site mutations engineered in *Mm*PylRS or *Mb*PylRS to members of the ΔNPylRS class, such as *Ma*PylRS^33^.

While transfer of active site mutations into the ΔNPylRS class has, in some cases, produced variants with incorporation efficiencies comparable to that of the PylRS in which those mutations were evolved^17,34,35^, in other cases, transplanting the active site mutations resulted in enzymes with poor activity, necessitating directed evolution^33,36^. Nevertheless, we decided to explore active site transplantation as an approach to engineer *Ma*PylRS to incorporate photocaged ncAAs, since no *Ma*PylRS variants currently exist for incorporating photocaged variants of canonical amino acids.

The photocaged tyrosines NPY, ONBY, and MNPY have been incorporated using the engineered *Mm*PylRS variant *Mm*NPYRS^37^, the photocaged lysines PCK and HCK have been incorporated using engineered *Mm*PylRS variants *Mm*PCKRS^29,38^ and *Mm*HCKRS^38,39^. The photocaged cysteine PCC2 has been incorporated using the engineered *Mm*PylRS variant *Mm*PCC2RS^40^. We have previously used these variants in *C. elegans*, for efficient incorporation of NPY and ONBY^41^, PCK^26,38^, HCK^38^, and PCC2^25^.

To generate candidate variants of *Ma*PylRS to incorporate photocaged ncAAs, we first aligned the sequences of *Mm*PylRS and *Ma*PylRS and identified residues in *Ma*PylRS corresponding to residues mutated in each *Mm*PylRS variant (**Fig. 5a**). For each *Mm*PylRS variant, we constructed three rounds of *Ma*PylRS variants. The first round of variants was constructed by direct transplantation of the respective mutations, resulting in *Ma*NPYRS1, *Ma*PCKRS1, *Ma*HCKRS1, and *Ma*PCC2RS1. The second round of variants was constructed by additionally introducing the mutations H227I and Y228P, which have previously been shown to improve incorporation efficiency of some ncAAs by *Ma*PylRS^42^, resulting in *Ma*NPYRS2, *Ma*PCKRS2, *Ma*HCKRS2, and *Ma*PCC2RS2. For the third round of variants, we introduced further mutations V42I, K181E, and A190V into the second round of variants. These mutations were originally developed by directed evolution and are reported to result in a hyperactive variant of *Ma*PylRS^43^. We named these variants *Ma*NPYRS3, *Ma*PCKRS3, *Ma*HCKRS3, and *Ma*PCC2RS3.

**Figure 5.**
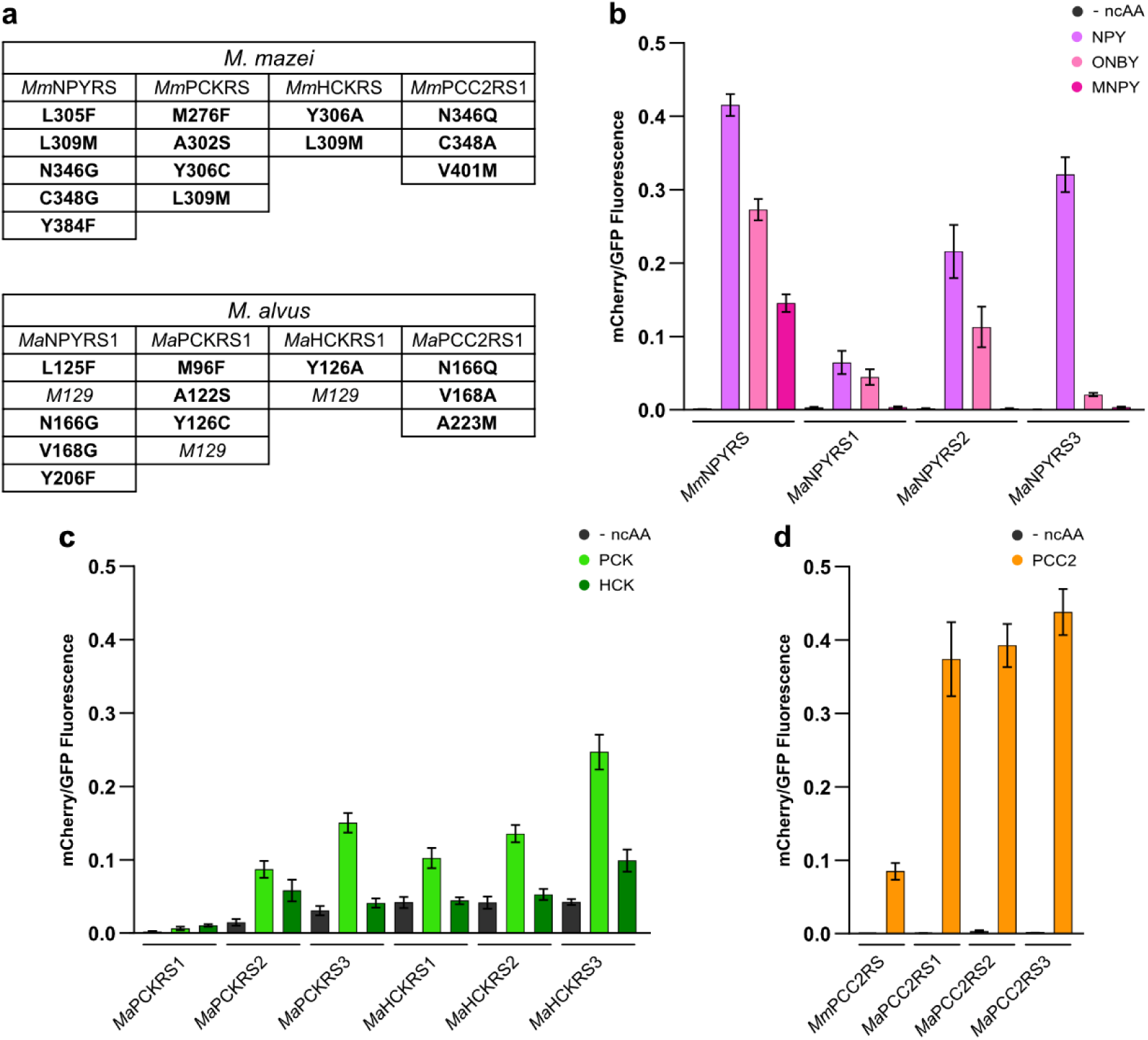
Engineering *Ma*PylRS to incorporate Photocaged ncAAs. **a** The top panel summarises the mutations in *Mm*PylRS that define the variants used to incorporate photocaged ncAAs. The bottom panel shows the mutations made at the homologous residues in *Ma*PylRS to generate the first round of candidate variants to incorporate photocaged ncAAs. Note that L309 in *Mm*PylRS corresponds to M129 in *Ma*PylRS and therefore remains unchanged. **b** mCherry to GFP fluorescence ratio of worms expressing either *Mm*NPYRS or an engineered *Ma*PylRS variant grown in the presence of NPY, ONBY, MNPY, or in the absence of ncAA. Error bars represent the SEM. **c** mCherry to GFP fluorescence ratio of worms expressing engineered *Ma*PylRS variants based on *Mm*PCKRS and *Mm*HCKRS, grown in the presence of PCK, HCK, or in the absence of ncAA. Error bars represent the SEM. **d** mCherry to GFP fluorescence ratio of worms expressing either *Mm*PCC2RS or an engineered *Ma*PylRS variant grown in the presence or absence of PCC2. Error bars represent the SEM.

To test these candidate variants, we generated transgenic strains expressing each *Ma*PylRS variant along with tRNA^Ma10^ and the GFP::TAG::mCherry reporter construct. We grew transgenic animals from the L4 larval stage for 24 hours on NGM agar plates supplemented with 1 mM of their respective ncAA, or in the absence of ncAA, and then assayed them for GFP and mCherry fluorescence. For comparison, we generated and assayed equivalent strains expressing each of the respective *Mm*PylRS variants together with tRNA^M15^.

We found that *Ma*NPYRS1 yielded only low incorporation efficiencies compared to *Mm*NPYRS, reaching only 15.2%, 16.3%, and 1.3% of the incorporation efficiency of *Mm*NPYRS for NPY, ONBY, and MNPY, respectively (**Fig. 5b**). *Ma*NPYRS2 significantly improved on *Ma*NPYRS1, reaching 51.9% and 41.4% of *Mm*NPYRS levels for NPY and ONBY, respectively. *Ma*NPYRS3 showed further improvement in incorporation efficiency with NPY, reaching 77.2% of *Mm*NPYRS. Interestingly, this enzyme had decreased activity with ONBY, suggesting that these mutations, which lie outside the active site, can differentially affect activity depending on the substrate. Similar to *Ma*NPYRS1, both *Ma*NPYRS2 and *Ma*NPYRS3 showed only negligible activity towards MNPY.

The *Mm*PylRS variants *Mm*PCKRS^29^ and *Mm*HCKRS^39^ were evolved separately using the photocaged lysines PCK and HCK, respectively. However, we previously found that the enzymes show significant cross-reactivity towards the alternative photocaged lysine^38^. We therefore tested all *Ma*PCKRS and *Ma*HCKRS variants with both PCK and HCK. For both photocaged lysines, *Ma*PCKRS1 yielded only low levels of incorporation (**Fig. 5c**). These were improved in *Ma*PCKRS2, and even further in *Ma*PCKRS3, however at the cost of increased background in the absence of ncAA. All *Ma*HCKRS variants showed high levels of background in the absence of ncAA. However, each consecutive variant did result in improved efficiency for both PCK and HCK. *Ma*HCKRS3 yielded the highest incorporation efficiencies of all *Ma*PCKRS and *Ma*HCKRS variants for both PCK and HCK.

Interestingly, when we tested incorporation of the photocaged cysteine PCC2, all *Ma*PCC2RS variants significantly outperformed the parental *Mm*PCC2RS2, with *Ma*PCC2RS1 already improving incorporation 4.4-fold (**Fig. 5d**). The subsequent *Ma*PCC2RS2 and *Ma*PCC2RS3 variants only provided marginal non-significant improvements to *Ma*PCC2RS1, and *Ma*PCC2RS2 displayed weak but clearly detectable background in the absence of PCC2.

For further experiments we focussed on the following *Ma*PylRS and ncAA combinations: *Ma*NPYRS2 for incorporation of NPY and ONBY, *Ma*NPYRS3 for NPY, and MaPCC2RS1 for PCC2.

### Optical control of FLP and Cre using engineered PylRS variants

Having established double ncAA incorporation in *C. elegans* and having engineered variants of *Ma*PylRS capable of efficiently incorporating photocaged ncAAs, we went on to apply our newly developed system.

As a first application, we decided on the optical control of the site-specific DNA recombinases (SSRs) FLP and Cre, which are both widely used, separately and in combination, to control gene expression in a variety of systems. They have previously been photocaged by us and others, using *Mm*PylRS variants to replace lysine, tyrosine, or cysteine residues in their active site with photocaged variants^26,38,44–46^. The use of two different photocaged ncAAs, incorporated by mutually orthogonal PylRS/tRNA^Pyl^ systems would allow the activity of FLP and Cre to be independently photocaged in the same animal, enabling separate optical control of two target genes.

To find the most efficient PylRS variant and ncAA to photocage each SSR, we first determined the efficiency of photocaged FLP (PC-FLP) and photocaged Cre (PC-Cre) when expressed by each engineered *Ma*PylRS variant, compared to the respective *Mm*PylRS variants. We generated transgenic strains containing three genetic components: an *Ma*PylRS/tRNA^Ma10^ or *Mm*PylRS/tRNA^M15^ pair engineered for incorporating a specific photocaged ncAA, a FLP or Cre construct containing a TAG amber stop codon at the desired amino acid position, and a Citrine2 fluorescent reporter gene whose expression is dependent upon activated FLP or Cre (**Fig. 6a, b**). To visualise cells containing the transgenes, we also included a constitutively expressed red mKate2 reporter gene in an artificial operon with FLP and Cre.

**Figure 6.**
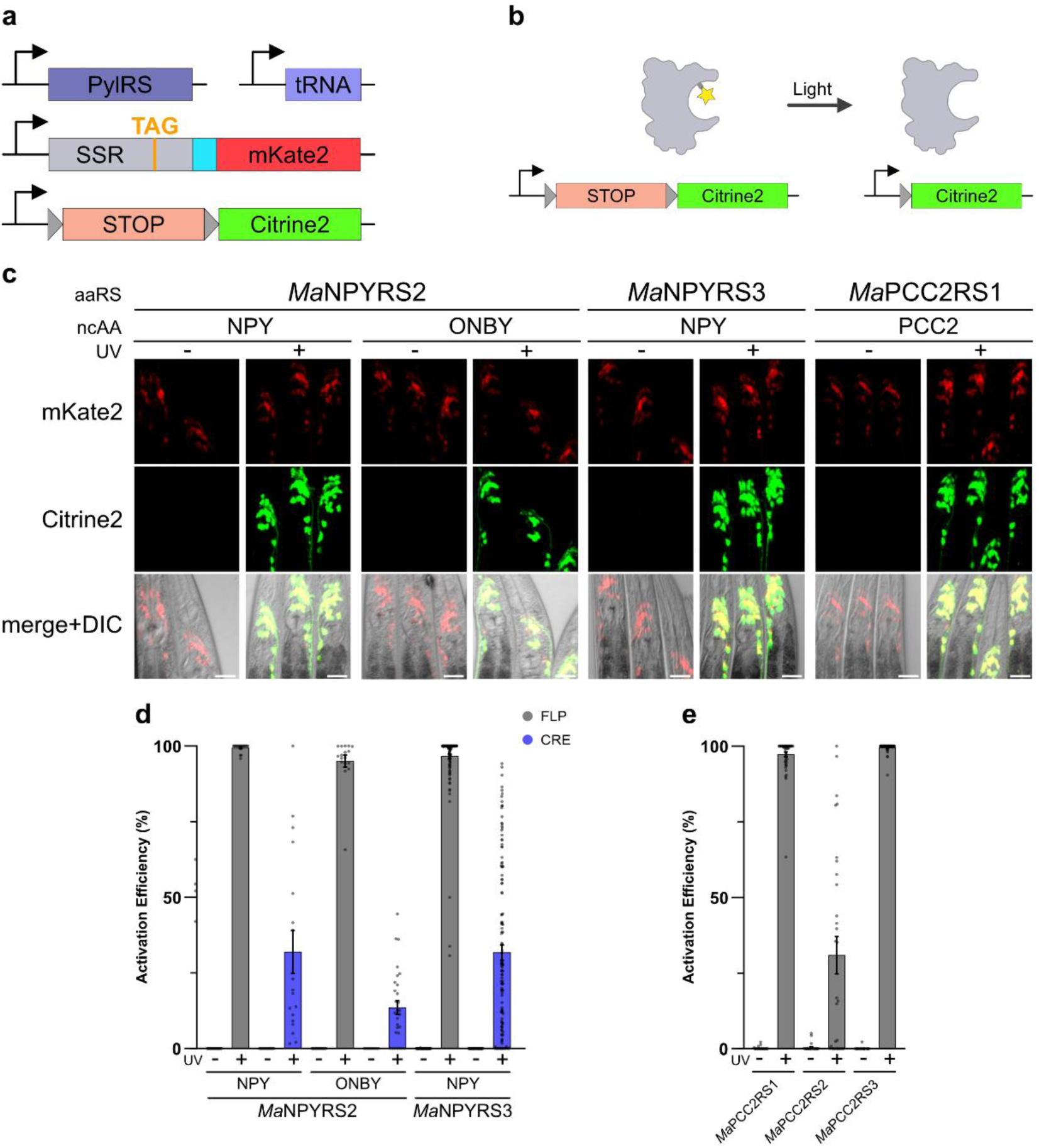
Photocaging FLP and Cre using engineered *Ma*PylRS variants. **a** Constructs transformed into *C. elegans* to express PC-FLP and PC-Cre and visualise their activity. The SSR gene in grey represents FLP or Cre. **b** Schematic depicting expression of the Citrine2 reporter following activation of a photocaged recombinase. The photocaged recombinase, which contains a photocaged ncAA in its active site, is unable to excise the transcriptional terminator inserted between the Citrine2 reporter gene and its promoter. Following illumination, the photocaging group within the photocaged recombinase is cleaved, producing active recombinase, which is able to excise the transcriptional terminator, allowing Citrine2 expression. **c** Fluorescence microscopy images of the head regions of *C. elegans* using select *Ma*PylRS variants to express PC-FLP. Scale bars represent 20 μm. **d** Activation efficiency of PC-FLP and PC-Cre using photocaged variants of tyrosine expressed by *Ma*NPYRS2, or *Ma*NPYRS3. Dots represent individual animals assayed. Error bars represent the SEM. **e** Activation efficiency of PC-FLP(C189PCC2) expressed by either of the three engineered *Ma*PCC2RS variants. Dots represent individual animals assayed. Error bars represent the SEM.

For photocaging, we selected residues that have successfully been photocaged previously using the *Mm*PylRS system. For FLP and Cre, we chose the catalytic tyrosines Y343 and Y324, respectively^38,44^. For FLP, we also included an essential active site cysteine, C189^38,47^, which is not present in Cre.

To incorporate photocaged variants of tyrosine into FLP and Cre, we used the *Ma*NPYRS2/tRNA^Ma10^ pair to direct incorporation of NPY and ONBY, the *Ma*NPYRS3/tRNA^Ma10^ pair to direct incorporation of NPY, as well as the established *Mm*NPYRS/tRNA^M15^ pair to direct incorporation of NPY, ONBY, and MNPY. To incorporate PCC2 in lieu of FLP C189, we used the three *Ma*PCC2RS variants, as well as *Mm*PCC2RS.

To characterise the activity of PC-FLP and PC-Cre expressed using either engineered *Ma*PylRS or *Mm*PylRS variants, we grew transgenic animals on 1 mM of the respective ncAA from the L1 larval stage for 24 hours and then illuminated them with 365 nm light to uncage the ncAA and thus activate the SSR^38^. 24 hours after illumination we assayed for Citrine2 fluorescence as a measure of SSR activity and found clearly visible Citrine2 expression for all PylRS - SSR - ncAA combinations (**Fig. 6c**).

When comparing target gene activation efficiencies, we found that PC-FLP(Y343ONBY) expressed using *Ma*NPYRS2 and PC-FLP(Y343NPY) expressed using *Ma*NPYRS2 or *Ma*NPYRS3 resulted in near 100% activation efficiency (**Fig. 6d**). When testing PC-Cre however, we found that PC-Cre(Y324NPY) expressed using *Ma*NPYRS2 or *Ma*NPYRS3 resulted in lower activation efficiencies of 46.4% and 46.1% respectively. Similarly, PC-Cre(Y324ONBY) expressed using *Ma*NPYRS2 resulted in an activation efficiency of only 13.6%. We could not detect any Citrine2 in animals grown on ncAA but not uncaged (**Fig. 6d**).

For PC-FLP(C189PCC2), quantifying the activation efficiency revealed that *Ma*PCC2RS1 and *Ma*PCC2RS3 variants both provided near 100% efficiency, while the *Ma*PCC2RS2 variant yielded an activation efficiency of 30.9% (**Fig. 6e**).

As *Ma*PCKRS3 and *Ma*HCKRS3 were the most efficient amongst *Ma*PylRS variants that incorporate the photocaged lysines PCK and HCK (**Fig. 5c**), we tested both for expression of PC-FLP(K223) and PC-Cre(K201). However, neither proved effective and we did not see expression of the Citrine2 reporter in any of the strains we assayed after growth on ncAA and uncaging.

### Co-expression of PC-FLP and PC-Cre using two photocaged ncAAs and optical control using two wavelengths

Having found that the newly engineered variants of *Ma*PylRS can be used to express variants of PC-FLP and PC-Cre, and that in some cases these are activated at least as efficiently as when expressed using the parental *Mm*PylRS variants, we next sought to develop a system using *M*mPylRS and *Ma*PylRS variants together to express PC-FLP and PC-Cre photocaged by different ncAAs in the same animal.

Such a system would allow us to select which target gene to express by choosing which ncAA to feed to the animals, and therefore which photocaged SSR (PC-SSR) to express. Additionally, since photocaged ncAAs can differ in their absorption spectra^38^, we reasoned that it may be possible to express SSRs that can be selectively activated using different wavelengths.

An important consideration in designing such a system is therefore the activation spectrum of each variant of PC-FLP and PC-Cre to be expressed. Previously, we have shown that different photocaged ncAAs imbue PC-FLP with different activation properties, with some resulting in PC-FLP variants that can be activated at longer wavelengths than others^38^. For example, PC-FLP(C189PCC2) can be efficiently activated at 435 nm, whereas PC-FLP(Y343NPY) cannot^38^. In principle, this property offers the possibility of creating a system in which PC-FLP(C189PCC2) is activated at 435 nm, while PC-Cre is activated only at 365 nm, giving greater flexibility for optically controlling gene expression and potentially allowing independent activation even when PC-FLP and PC-Cre are expressed simultaneously.

We indeed found that PC-FLP(C189PCC2) expressed using *Ma*PCC2RS1 and illuminated at either 365 nm or 435 nm was activated with near 100% efficiency at both wavelengths, while in contrast PC-Cre(Y324NPY) expressed using *Mm*NPYRS could only be activated using 365nm but showed no activation at 435nm. (**Fig. 7a**).

**Figure 7.**
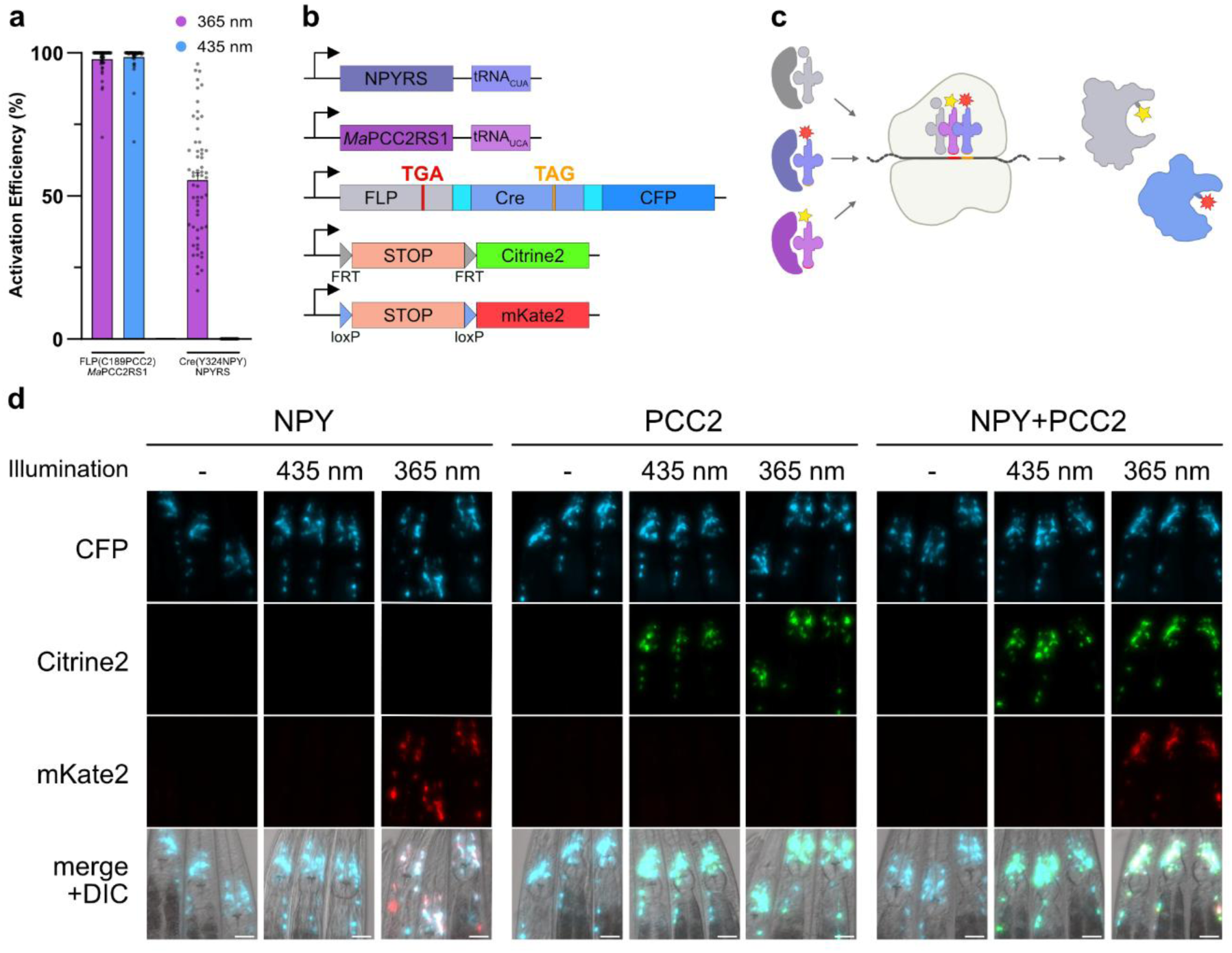
Expression of PC-FLP and PC-Cre in a single animal using two ncAAs. **a** Activation efficiency at 365 nm and 435 nm of PC-FLP(C189PCC2) expressed by *Ma*PCC2RS1, and of PC-Cre(Y324NPY) expressed by *Mm*NPYRS. Dots represent individual animals assayed. Error bars represent the SEM. **b** Constructs transformed into *C. elegans* to express PC-FLP and PC-Cre and to visualise their activity. **c** Schematic depicting the expression of PC-FLP and PC-Cre using two different photocaged ncAAs. **d** Fluorescence microscopy images of the head regions of *C. elegans* grown in the presence of NPY alone, PCC2 alone, or both NPY and PCC2 together to express PC-Cre or PC-FLP respectively. Animals were illuminated at either 365 nm, 435 nm, or kept in the dark. Scale bars represent 20 μm.

To use PC-FLP(C189PCC2) and PC-Cre(Y324NPY) in the same animal, we generated transgenic strains by co-transforming *Ma*PCC2RS1/tRNA^Ma10^, *Mm*NPYRS/tRNA^M15^ , FLP(C189TGA) and Cre(Y324TAG) (**Fig. 7b, c**). The FLP and Cre genes were expressed in an artificial operon that also included a constitutively expressed CFP for visualisation. We furthermore co-transformed two fluorescent reporter constructs to assay recombination. One reporter consisted of a Citrine2 gene that was separated from its promoter by a transcriptional terminator flanked by FRT sites, and the other reporter consisted of an mKate2 gene that was separated from its promoter by a transcriptional terminator flanked by loxP sites. FLP activity would thus result in Citrine2 expression, and Cre activity would result in mKate2 expression.

We first tested whether we could determine which target gene to express through the ncAA we provided. We grew transgenic worms from the L1 larval stage for 24 hours on NGM agar plates containing either PCC2 or NPY and assayed them for Citrine2 and mKate2 expression 24h after uncaging.

We found that by feeding NPY alone, to induce expression of PC-Cre(324NPY), followed by illumination with 365 nm light, we could switch on expression of mKate2 while leaving Citrine2 switched off. Conversely feeding PCC2 to induce expression of PC-FLP(C189PCC2), followed by illumination with 365nm switched on expression of Citrine2 but did not switch on mKate2. For both cases, we saw no target gene expression without illumination. This demonstrated that the two PylRS - SSR - ncAA systems were fully orthogonal and that it was indeed possible to control expression of a specific target gene through selectively feeding NPY or PCC2, followed by uncaging to activate the SSR.

We then tested the responsiveness to 365nm vs 435nm. As expected, 435nm light had no effect in animals expressing PC-Cre(Y324NPY) and we saw no mKate 2 expression, similar to the non-illuminated control. This was in contrast to animals grown on PCC2 and therefore expressing PC-FLP(C189PCC2). Here, illumination with either 365nm or 435nm switched on expression of Citrine2. We then went on and grew the animals in the presence of both NPY and PCC2 and found that when both photo-caged SSRs were expressed together it was also possible to activate PC-FLP(C189PCC2) alone through illumination with 435nm, without affecting PC-Cre(Y324NPY), and to selectively switch on only Citrine2. Illumination with 365nm on the other hand, as expected led to activation of both SSRs, resulting in expression of both target genes Citrine2 and mKate2 together (**Fig. 7d**).

We thus combined our system for double incorporation of ncAA, our engineered *M. alvus* PylRS variants for incorporation of photocaged ncAAs, and our previously established PC-FLP and PC-Cre variants to develop a method for the optical control of two target genes within *C. elegans*. This method allows us to control which SSR becomes active, and therefore which target gene is switched on, either based on which ncAA is provided, or which activating wavelength is used, thus providing flexible control over the expression of two genes in an animal.

## DISCUSSION

In this paper, we establish the simultaneous and independent incorporation of two ncAAs for the first time in an animal. The ability to incorporate two ncAAs in a multicellular organism provides the basis for novel experimental approaches that are otherwise inaccessible. We demonstrate this by applying the double ncAA incorporation system to independently photocage the activity FLP and Cre within the same animal, enabling modular optical control over the expression of two target genes.

To incorporate two ncAAs, we first established the PylRS/tRNA^Pyl^ pair from *M. alvus*. in *C. elegans* and demonstrated its suitability for GCE as well as its mutual orthogonality with the previously established *M. mazei* PylRS/tRNA pair in the worm. This suggests the possibility that these two PylRS/tRNA^Pyl^ pairs may also be applicable to incorporate two ncAAs in the other animal models in which genetic code expansion has been established^48^. By extension, the transferability of the *M. alvus* PylRS/tRNA^Pyl^ pair used here suggests that it may be possible to also apply, in multicellular organisms, additional mutually orthogonal PylRS/tRNA^Pyl^ pairs that have been developed as part of triply^18^, quadruply^49^, or quintuply^19^ orthogonal groups, eventually leading to systems that go beyond the simultaneous incorporation of two ncAA in multicellular organisms.

The new quadruplet codons established here for use in *C. elegans* in turn provide a potential path towards the additional coding space required to incorporate multiple ncAAs using such additional mutually orthogonal pairs. The successful transplanting of several optimised anticodon loops optimised for quadruplet decoding onto a eukaryote-optimised *Mm*tRNA scaffold to create hybrid tRNA variants, suggests it may be possible to apply this approach to create additional quadruplet decoding tRNAs, for example hybrid variants of *Ma*tRNA^Pyl^ or tRNAs from other mutually orthogonal pairs.

The expansion of a metazoan genetic code beyond a single ncAA allowing the concurrent incorporation of two ncAAs further augments the GCE toolbox and will enable the development of new experimental tools and approaches. In bacteria and mammalian cell culture, double ncAA incorporation has been used in applications such as to site-specifically attach a drug and fluorescent label to an antibody^50^, to covalently link proteins^21^, or to attach distinct fluorophores to visualise separate proteins or to act as FRET probes^11^. The development of a system that can form the basis for adapting these tools to metazoan systems opens up exciting new methodological opportunities. As the first example for an application, we establish dual photocaged SSRs, which are particularly suited for multicellular organisms, where precise targeting of gene expression is challenging and where photocaged SSRs allow control down to single-cell resolution^26,38^.

Future work to validate the transferability into other multicellular organisms of the mutually orthogonal PylRS/tRNA^Pyl^ pair and blank codons established here may provide the basis for developing tools that can be applied across model organisms, facilitating tool development in other genetically amenable models.

To engineer the ncAA specificity of the orthogonal *Ma*PylRS/tRNA^Pyl^ to incorporate photocaged amino acids, we began by direct transplantation of active site residues. This approach was successful at producing highly active *Ma*PylRS variants for incorporating PCC2 but did not immediately yield variants for incorporating photocaged variants of tyrosine or lysine. In the case of tyrosine, incorporation efficiency towards two photocaged variants was improved by introducing additional mutations that were previously shown to provide general improvements to incorporation efficiency. This demonstrates a potential workflow for rapidly generating and improving *Ma*PylRS variants for incorporating desired ncAAs. Since these experiments were performed, new research has been published describing additional mutations which appear to also generally improve incorporation efficiency^51,52^, and which may therefore serve in this workflow. In the case of *Ma*PylRS variants for incorporating photocaged variants of lysine, we found all to have low incorporation efficiency and high levels of background. Neither of these was fully resolved by the addition of additional mutations, highlighting a case where active site transplantation was not successful and directed evolution may be necessary.

Combining our newly engineered *Ma*PylRS variants with the established *Mm*PylRS system allowed us to express the photocaged SSRs PC-FLP and PC-Cre in a single animal to control the expression of two target genes. Crucially, we took advantage of the distinct uncaging properties between PCC2 used to photocage FLP and NPY used to photocage Cre, which made it possible to spectrally separate PC-FLP activation from PC-Cre activation, introducing selective control over gene expression based on the ncAA and wavelength provided. We show that this system can be applied to optically control the expression of two genes in *C. elegans*. Importantly, since the activating light can be precisely targeted, it will be possible to carry out such studies within specific organs or cells in multicellular organisms. Furthermore, PC-FLP can be activated separately from PC-Cre to express its respective target gene, a capability that could aid studies benefitting from sequential control of gene expression, for example to decode gene regulatory networks. While this work has been carried out in *C. elegans*, FLP, Cre, and genetic code expansion have been established in other animal models, such as zebrafish, *Drosophila*, and mice, suggesting that this work may also be transferable to those systems.

The system we have developed here can be applied to control expression of two genes based on the ncAA and the wavelength provided. Taken together, these two inputs can be used to output the four possible gene expression conditions. Alternative systems for optical control of gene expression exist, based on split SSRs that are reconstituted using photosensitive protein domains, which have been used to independently control the activity of Cre and FLP using red and blue light illumination^53^. These however can require extended illumination times, compared to PC-FLP/Cre which can be irreversibly activated by illumination in the low millisecond range^26,38^. In addition, photocaged SSRs generally do not show background activity and are not activated by wavelengths used for other optogenetic approaches or imaging.

Furthermore, the photocaged SSR system has the potential to be extended beyond just two recombinases. SSRs in the tyrosine site-specific recombinase family to which FLP and Cre belong possess highly conserved active site residues^54^. Given that the approach of replacing active site residues with photocaged counterparts has been successful for both FLP and Cre, it is likely that it will be transferable to other SSRs. Thus far, the activity of FLP has been photocaged using photocaged versions of tyrosine, lysine, and cysteine^38^, while the activity of Cre has been photocaged using photocaged versions of tyrosine^44^, lysine^26,45,46^, and histidine^55^, and the same ncAAs might be adapted to develop additional photocaged SSRs. The existence of mutually orthogonal translation systems could make it possible to co-express two or more such photocaged SSRs, each activated using different illumination conditions, such as UV, blue light, green light^56^, or near infrared light^39^ depending on the photocaged ncAA used. Additionally, newly developed photocaged ncAA variants could be easily integrated into the existing system to upgrade the PC-SSRs without the need for additional engineering. Additional photocaged SSRs could also be applied to complement systems using optically controlled split FLP and Cre and activated using blue and red light, in order to optically and independently control the expression of more than two genes.

## METHODS

### C. elegans strains

Strains were maintained on nematode growth medium (NGM) agar plates under standard conditions unless otherwise indicated^57,58^. Transgenic strains were generated by biolistic bombardment using hygromycin B as a selection marker^23,26,59^. Strains were maintained on Nematode growth medium (NGM) supplemented with hygromycin at a concentration of 0.3 mg/ml (Formedium). Strains were constructed in either the N2 (wild type) or *smg-6(ok1794)* genetic background. The *smg-6(ok1794)* background, which is deficient in nonsense mediated decay, was used for all strain expressing the GFP::mCherry fluorescent reporter used to measure direct incorporation efficiencies in order to stabilise reporter mRNA. All strain expressing PC-FLP or PC-Cre were constructed in the N2 background. Transgenic strains are described in Supplementary Table 1.

### Plasmid construction

Expression plasmids transformed into *C. elegans* were assembled from entry plasmids and destination vectors using the Gateway LR Clonase II Plus Enzyme Mix (Thermo Fisher Scientific).

DNA fragments and oligonucleotides used in the generation of entry plasmids were custom synthesised by IDT. Description and construction of expression plasmids is detailed in Supplementary Table 2. Entry plasmids used in the LR reactions are described in Supplementary table 3. The sequences for the genes described in this work are given in Supplementary Note 1.

### ncAA feeding

PCK^29^ and PCC2^40^ were custom synthesised by ChiroBlock GmbH, Germany. HCK^39^ was custom synthesised by Inochem, United Kingdom. NPY^37^ and MNPY^37^ were custom synthesised by NewChem Technologies, UK. ONBY^60^ was purchased from Fluorochem (F447104).

NGM agar plates containing ncAA were prepared by dissolving the ncAA in 0.1 ml of HCl or NaOH and then adding it to the molten NGM agar. The HCl or NaOH were then neutralised by adding equimolar amounts of NaOH or HCl respectively to the molten NGM ncAA solution. Approximately 1.5ml of NGM agar was used per 35mm plate. PCK was dissolved in 0.2 M HCl, HCK was dissolved in 0.1 M NaOH, NPY was dissolved in 1 M NaOH, MNPY was dissolved in 0.1 M NaOH, ONBY was dissolved in 1 M NaOH, PCC2 was dissolved in 0.5 M HCl.

Animals were age-synchronised by settling and added to NGM ncAA plates as L1 larvae unless otherwise stated. Freeze-dried OP50 (LabTIE) reconstituted according to the manufacturer’s instructions was added for food. Animals were grown on the NGM ncAA plates for 24 hours unless otherwise stated.

### Western blotting

Worms were grown on NGM plates supplemented with ncAA for 24 hours and then washed off using M9 buffer containing 0.001% Triton X-100. The worms were settled and the supernatant was removed and replaced with twice the pellet volume of a lysis buffer composed of a 4:1 ratio of 4×LDS loading buffer (Thermo Fisher Scientific) and NuPAGE Sample Reducing Agent (Thermo Fisher Scientific).

Worms were then frozen at -20 °C overnight and then incubated for 15 minutes at 95 °C. Samples were run on precast Bolt 4-12% gels (Thermo Fisher Scientific) at 200 V for 20 minutes and then transferred onto a nitrocellulose membrane using and iBlot2 (Thermo Fisher Scientific). After transfer, membranes were blocked for 1 h at room temperature using PBST (PBS + 0.1% Tween-20) supplemented with 5% milk powder. Incubation with primary antibodies was carried out at 4°C overnight. Blots were washed 6 times for 5 min in PBST + 5% milk powder. Blots were then incubated with secondary antibodies diluted in PBST + 5% milk powder for 1 h at room temperature, followed by three washes of 5 min using PBST + 5% milk powder and one wash with PBS.

Primary antibodies used were mouse anti-GFP (clones 7.1 and 13.1, Roche) and Rat anti-HA (clone 3F10) (Roche), at dilutions of 1:4000 and 1:2000 respectively. Secondary antibodies used were horse anti-mouse IgG HRP (Cell Signalling Technology) and goat anti-rat IgG HRP (Thermo Fisher Scientific), both at a dilution of 1:5000. The HRP substrates ECL Western (Thermo Fisher Scientific) and SuperSignal West Femto (Thermo Fisher Scientific) were used, and detection was carried out using photographic film.

### Photocaged Recombinase activation

Worms were grown on NGM agar plates supplemented with ncAA as described and washed off with M9 supplemented with 0.001% Triton-X100 to prevent animals sticking to pipette tips. To test activation, animals were illuminated in a CL-1000L crosslinker at 365 nm (Analytik Jena) for 5 minutes. To test activation at defined wavelengths, a pE-4000 light source (CoolLED) was used as previously described^38^. Worms were illuminated with the 365 nm and 435 nm LEDs at 100% power for 5 minutes, equivalent to 30 mW and 170 mW respectively.

### Imaging and analysis

All imaging was carried out on a Zeiss Axio Imager M2 using ZEN software (Zeiss). Worms were anaesthetized with 50 mM NaN_3_ and mounted on 2.5% agar pads on microscope slides. All images compared were taken at the same magnification and exposure settings. All image analysis was carried out on Fiji version 2.14.0.

To score the mCherry/GFP fluorescence ratio in animals expressing the GFP::mCherry ncAA incorporation reporter, fluorescence z-stack images of worms were acquired, with each image containing multiple individuals. For each channel, a single 2D image was created from the z-stack image by sum intensity projection. Regions where the GFP fluorescent marker was expressed were identified using Huang thresholding^61^ and used to generated a mask. The mask was then separately applied to the GFP and the mCherry channels. In the mCherry channel, pixels within the mask that did not exceed a signal threshold defined equally for all images were discarded. Fluorescence intensity histograms were then generated for both the masked GFP channel and the masked and thresholded mCherry channel. For each channel, a background value defined as the minimum pixel value in each channel prior to masking was subtracted from each histogram bin. The total fluorescence intensity for each channel was then calculated by summing the products of each histogram bin value and its corresponding count.

Quantification of the activation efficiency of PC-FLP and PC-Cre was carried out as previously described^38^. Briefly, sum projection images in the Citrine2 and mKate2 channels across the whole width of the worm’s head were used to compare the area in which they were expressed. The fluorescent mKate2 marker within worms was first automatically identified using Li thresholding^62^. This thresholding was used to create a mask that was applied onto the fluorescent Citrine2 reporter channel. Within this mask, the number of pixels that exceeded a signal threshold defined equally for all images were counted as signal. Activation efficiency was calculated as the ratio of the number of signal pixels to mask pixels, expressed as a percentage

## ACKNOWLEDGEMENTS

The work was supported by the European Research Council [ERC-StG-679990], the BBSRC [BB/Y006380/1 & BB/W014610/1], The Royal Society, The Muir Maxwell Epilepsy Centre. We thank members of the Greiss and Doitsidou labs for helpful suggestions.

## AUTHOR CONTRIBUTIONS

J.J.V.R. and S.G. conceived and designed experiments. J.J.V.R. performed experiments and analysed data. J.J.V.R. and SG wrote the manuscript.

