## Supplementary for "Expanding the Genetic Code of an Animal with Two Non-Canonical Amino Acids"

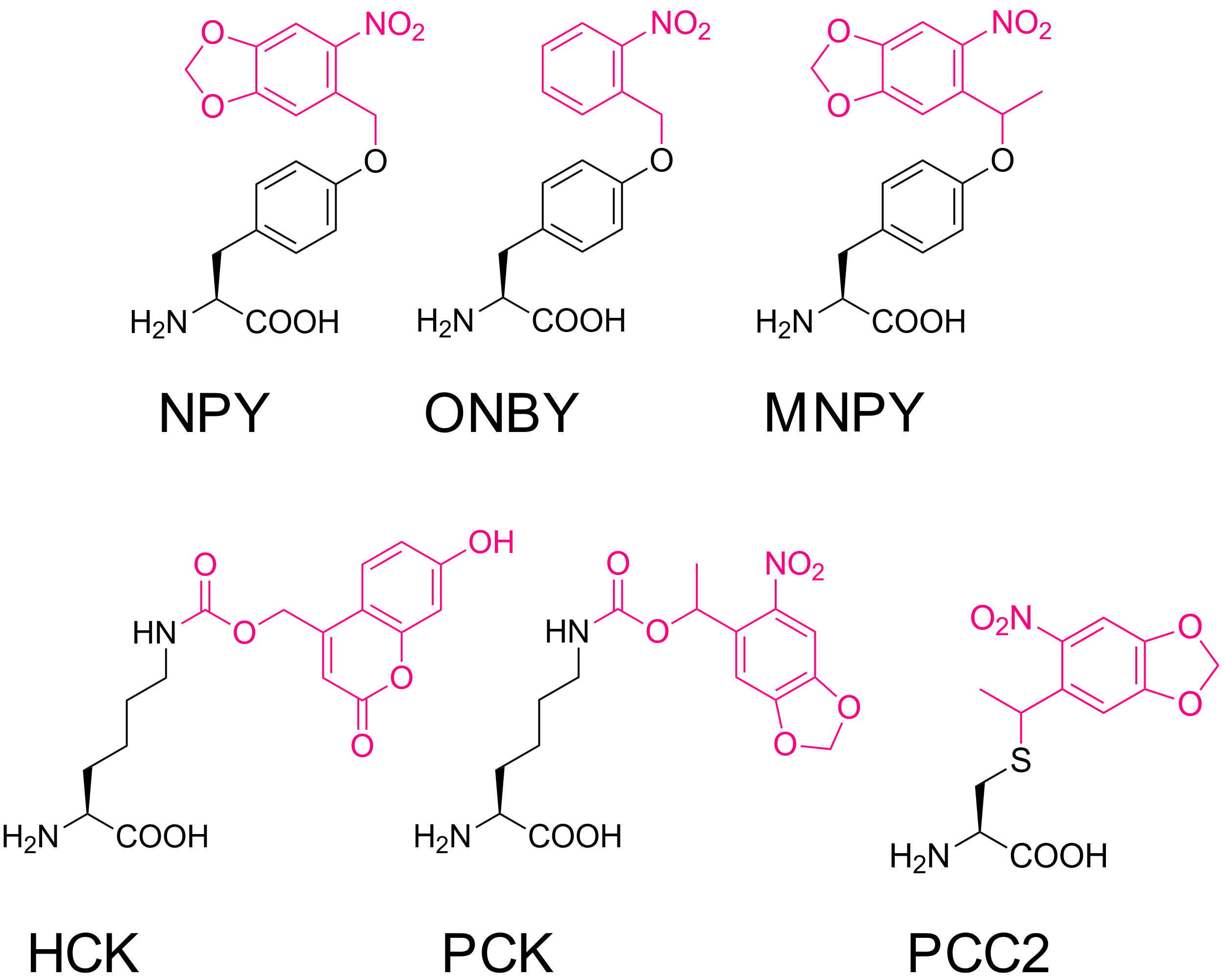

**Supplementary Fig. 1 Photocaged ncAAs used in this study.**

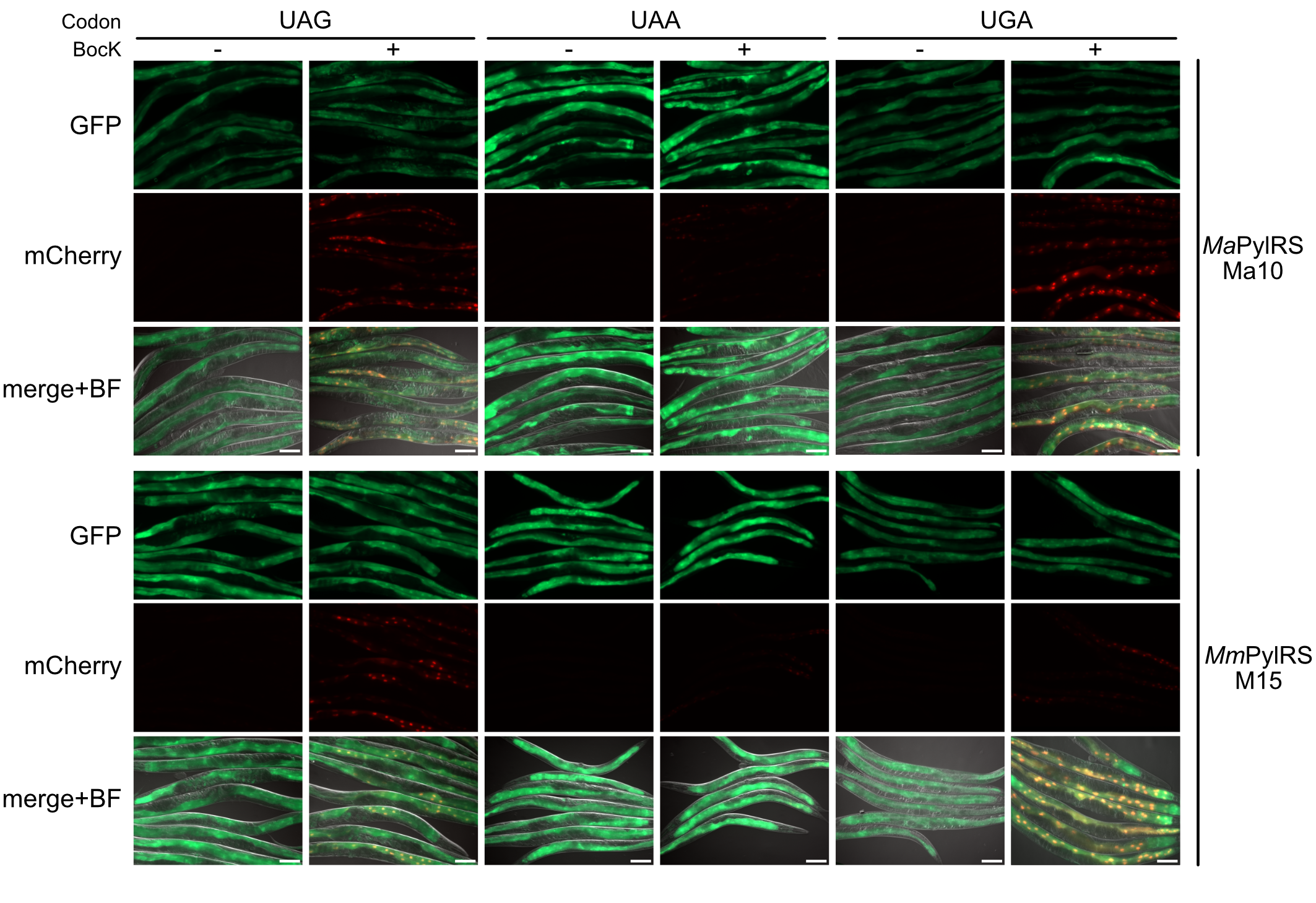

**Supplementary Fig. 2 Incorporation at the three stop codons.** Fluorescence microscopy images of worms expressing the GFP::mCherry reporter bearing either of the three stop codons in between the GFP and mCherry genes together with the *Ma*PylRS/^Ma10^tRNA or *Mm*PylRS/^M15^tRNA pair bearing the respective anticodon to decode the codon in the reporter. Scale bars represent 100 μm.

**Supplementary Table 1 Transgenic *C. elegans* strains.**

| Name | Description | Genetic Background | Expression Plasmids (ug) | Location in article |
| --- | --- | --- | --- | --- |
| SGR209 | *greEx190[sur-5p::HuSmad-4_NLS::MmPylRS::let-858_3’UTR; sur-5p::HygR::unc-54_3’UTR; rpr-1p::tRNA(M15)::sup-7_3’UTR; rps-0p::GFP::TAG::mCherry::HA::EGL-13_NLS::unc-54_3’UTR]* | *smg-6(ok1794)* | JJ59(2), SE149(7), SG88(3) | Fig. 1b, c, Fig. 2b, Fig. 3a, Fig. 3d, Supplementary Fig. 2 |
| SGR210 | *greEx191[sur-5p::HuSmad-4_NLS::MmPylRS::let-858_3’UTR; sur-5p::HygR::unc-54_3’UTR; rpr-1p::tRNA(M15)::sup-7_3’UTR; rps-0p::GFP::TAG::mCherry::HA::EGL-13_NLS::unc-54_3’UTR]* | *smg-6(ok1794)* | JJ59(2), SE149(7), SG88(3) | Fig. 1c, Fig. 3d |
| SGR211 | *greEx192[sur-5p::HuSmad-4_NLS::MmPylRS::let-858_3’UTR; sur-5p::HygR::unc-54_3’UTR; rpr-1p::tRNA(C15)::sup-7_3’UTR; rps-0p::GFP::TAG::mCherry::HA::EGL-13_NLS::unc-54_3’UTR]* | *smg-6(ok1794)* | JJ59(2), SE150(7), SG88(3) | Fig. 2b |
| SGR212 | *greEx193[sur-5p::HuSmad-4_NLS::MmPylRS::let-858_3’UTR; sur-5p::HygR::unc-54_3’UTR; rpr-1p::tRNA(G1hyb)::sup-7_3’UTR; rps-0p::GFP::TAG::mCherry::HA::EGL-13_NLS::unc-54_3’UTR]* | *smg-6(ok1794)* | JJ59(2), JJ57(7), SG88(3) | Fig. 2b |
| SGR213 | *greEx194[sur-5p::HuSmad-4_NLS::MmPylRS::let-858_3’UTR; sur-5p::HygR::unc-54_3’UTR; rpr-1p::tRNA(Ma6)::sup-7_3’UTR; rps-0p::GFP::TAG::mCherry::HA::EGL-13_NLS::unc-54_3’UTR]* | *smg-6(ok1794)* | JJ59(2), SE297(7), SG88(3) | Fig. 2b |
| SGR214 | *greEx195[sur-5p::HuSmad-4_NLS::MmPylRS::let-858_3’UTR; sur-5p::HygR::unc-54_3’UTR; rpr-1p::tRNA(Ma10)::sup-7_3’UTR; rps-0p::GFP::TAG::mCherry::HA::EGL-13_NLS::unc-54_3’UTR]* | *smg-6(ok1794)* | JJ59(2), SE298(7), SG88(3) | Fig. 2b |
| SGR215 | *greEx196[sur-5p::MaPylRS::let-858_3’UTR; sur-5p::HygR::unc-54_3’UTR; rpr-1p::tRNA(Ma6)::sup-7_3’UTR; rps-0p::GFP::TAG::mCherry::HA::EGL-13_NLS::unc-54_3’UTR]* | *smg-6(ok1794)* | KB375(2), SE297(7), SG88(3) | Fig 1b, c, Fig. 2b |
| SGR216 | *greEx197[sur-5p::MaPylRS::let-858_3’UTR; sur-5p::HygR::unc-54_3’UTR; rpr-1p::tRNA(Ma6)::sup-7_3’UTR; rps-0p::GFP::TAG::mCherry::HA::EGL-13_NLS::unc-54_3’UTR]* | *smg-6(ok1794)* | KB375(2), SE297(7), SG88(3) | Fig 1c |
| SGR217 | *greEx198[sur-5p::MaPylRS::let-858_3’UTR; sur-5p::HygR::unc-54_3’UTR; rpr-1p::tRNA(Ma10)::sup-7_3’UTR; rps-0p::GFP::TAG::mCherry::HA::EGL-13_NLS::unc-54_3’UTR]* | *smg-6(ok1794)* | KB375(2), SE298(7), SG88(3) | Fig 1b, c, Fig. 2b, Fig. 3d |
| SGR218 | *greEx199[sur-5p::MaPylRS::let-858_3’UTR; sur-5p::HygR::unc-54_3’UTR; rpr-1p::tRNA(Ma10)::sup-7_3’UTR; rps-0p::GFP::TAG::mCherry::HA::EGL-13_NLS::unc-54_3’UTR]* | *smg-6(ok1794)* | KB375(2), SE298(7), SG88(3) | Fig 1c, Fig. 3d, Supplementary Fig. 2 |
| SGR219 | *greEx200[sur-5p::MaPylRS::let-858_3’UTR; sur-5p::HygR::unc-54_3’UTR; rpr-1p::tRNA(M15)::sup-7_3’UTR; rps-0p::GFP::TAG::mCherry::HA::EGL-13_NLS::unc-54_3’UTR]* | *smg-6(ok1794)* | KB375(2), SE149(7), SG88(3) | Fig. 2b |
| SGR220 | *greEx201[sur-5p::MaPylRS::let-858_3’UTR; sur-5p::HygR::unc-54_3’UTR; rpr-1p::tRNA(C15)::sup-7_3’UTR; rps-0p::GFP::TAG::mCherry::HA::EGL-13_NLS::unc-54_3’UTR]* | *smg-6(ok1794)* | KB375(2), SE150(7), SG88(3) | Fig. 2b |
| SGR221 | *greEx202[sur-5p::MaPylRS::let-858_3’UTR; sur-5p::HygR::unc-54_3’UTR; rpr-1p::tRNA(G1hyb)::sup-7_3’UTR; rps-0p::GFP::TAG::mCherry::HA::EGL-13_NLS::unc-54_3’UTR]* | *smg-6(ok1794)* | KB375(2), JJ57(7), SG88(3) | Fig. 2b |
| SGR222 | *greEx203[sur-5p::FLAG::G1PylRS::let-858_3’UTR; sur-5p::HygR::unc-54_3’UTR; rpr-1p::tRNA(G1hyb)::sup-7_3’UTR; rps-0p::GFP::TAG::mCherry::HA::EGL-13_NLS::unc-54_3’UTR]* | *smg-6(ok1794)* | JJ60(2), JJ57(7), SG88(3) | Fig 1b, c, Fig. 2b |
| SGR223 | *greEx204[sur-5p::FLAG::G1PylRS::let-858_3’UTR; sur-5p::HygR::unc-54_3’UTR; rpr-1p::tRNA(G1hyb)::sup-7_3’UTR; rps-0p::GFP::TAG::mCherry::HA::EGL-13_NLS::unc-54_3’UTR]* | *smg-6(ok1794)* | JJ60(2), JJ57(7), SG88(3) | Fig 1c |
| SGR224 | *greEx205[sur-5p::FLAG::G1PylRS::let-858_3’UTR; sur-5p::HygR::unc-54_3’UTR; rpr-1p::tRNA(Ma6)::sup-7_3’UTR; rps-0p::GFP::TAG::mCherry::HA::EGL-13_NLS::unc-54_3’UTR]* | *smg-6(ok1794)* | JJ60(2), SE297(7), SG88(3) | Fig. 2b |
| SGR225 | *greEx206[sur-5p::FLAG::G1PylRS::let-858_3’UTR; sur-5p::HygR::unc-54_3’UTR; rpr-1p::tRNA(Ma10)::sup-7_3’UTR; rps-0p::GFP::TAG::mCherry::HA::EGL-13_NLS::unc-54_3’UTR]* | *smg-6(ok1794)* | JJ60(2), SE298(7), SG88(3) | Fig. 2b |
| SGR226 | *greEx207[sur-5p::FLAG::G1PylRS::let-858_3’UTR; sur-5p::HygR::unc-54_3’UTR; rpr-1p::tRNA(M15)::sup-7_3’UTR; rps-0p::GFP::TAG::mCherry::HA::EGL-13_NLS::unc-54_3’UTR]* | *smg-6(ok1794)* | JJ60(2), SE149(7), SG88(3) | Fig. 2b |
| SGR227 | *greEx208[sur-5p::FLAG::G1PylRS::let-858_3’UTR; sur-5p::HygR::unc-54_3’UTR; rpr-1p::tRNA(C15)::sup-7_3’UTR; rps-0p::GFP::TAG::mCherry::HA::EGL-13_NLS::unc-54_3’UTR]* | *smg-6(ok1794)* | JJ60(2), SE150(7), SG88(3) | Fig. 2b |
| SGR228 | *greEx209[sur-5p::HuSmad-4_NLS::MmPylRS::let-858_3’UTR; sur-5p::HygR::unc-54_3’UTR; rpr-1p::tRNA(M15)[UCA]::sup-7_3’UTR; rps-0p::GFP::TGA::mCherry::HA::EGL-13_NLS::unc-54_3’UTR]* | *smg-6(ok1794)* | JJ59(2), KB337(7), JJ108(3) | Fig. 3a, Fig. 3d |
| SGR229 | *greEx210[sur-5p::HuSmad-4_NLS::MmPylRS::let-858_3’UTR; sur-5p::HygR::unc-54_3’UTR; rpr-1p::tRNA(M15)[UCA]::sup-7_3’UTR; rps-0p::GFP::TGA::mCherry::HA::EGL-13_NLS::unc-54_3’UTR]* | *smg-6(ok1794)* | JJ59(2), KB337(7), JJ108(3) | Fig. 3d, Supplementary Fig. 2 |
| SGR230 | *greEx211[sur-5p::HuSmad-4_NLS::MmPylRS::let-858_3’UTR; sur-5p::HygR::unc-54_3’UTR; rpr-1p::tRNA(M15)[UCA]::sup-7_3’UTR; rps-0p::GFP::TGA::mCherry::HA::EGL-13_NLS::unc-54_3’UTR]* | *smg-6(ok1794)* | JJ59(2), KB337(7), JJ108(3) | Fig. 3d, Supplementary Fig. 2 |
| SGR231 | *greEx212[sur-5p::HuSmad-4_NLS::MmPylRS::let-858_3’UTR; sur-5p::HygR::unc-54_3’UTR; rpr-1p::tRNA(M15)[UUA]::sup-7_3’UTR; rps-0p::GFP::TAA::mCherry::HA::EGL-13_NLS::unc-54_3’UTR]* | *smg-6(ok1794)* | JJ59(2), JJ268(7), JJ266(3) | Fig. 3a, Fig. 3d, Supplementary Fig. 2 |
| SGR232 | *greEx213[sur-5p::MaPylRS::let-858_3’UTR; sur-5p::HygR::unc-54_3’UTR; rpr-1p::tRNA(Ma10)[UCA]::sup-7_3’UTR; rps-0p::GFP::TGA::mCherry::HA::EGL-13_NLS::unc-54_3’UTR]* | *smg-6(ok1794)* | KB375(2), JJ269(7), JJ108(3) | Fig. 3d |
| SGR233 | *greEx214[sur-5p::MaPylRS::let-858_3’UTR; sur-5p::HygR::unc-54_3’UTR; rpr-1p::tRNA(Ma10)[UCA]::sup-7_3’UTR; rps-0p::GFP::TGA::mCherry::HA::EGL-13_NLS::unc-54_3’UTR]* | *smg-6(ok1794)* | KB375(2), JJ269(7), JJ108(3) | Fig. 3d |
| SGR234 | *greEx215[sur-5p::MaPylRS::let-858_3’UTR; sur-5p::HygR::unc-54_3’UTR; rpr-1p::tRNA(Ma10)[UUA]::sup-7_3’UTR; rps-0p::GFP::TAA::mCherry::HA::EGL-13_NLS::unc-54_3’UTR]* | *smg-6(ok1794)* | KB375(2), JJ267(7), JJ266(3) | Fig. 3d |
| SGR235 | *greEx216[sur-5p::MaPylRS::let-858_3’UTR; sur-5p::HygR::unc-54_3’UTR; rpr-1p::tRNA(Ma10)[UUA]::sup-7_3’UTR; rps-0p::GFP::TAA::mCherry::HA::EGL-13_NLS::unc-54_3’UTR]* | *smg-6(ok1794)* | KB375(2), JJ267(7), JJ266(3) | Fig. 3d, Supplementary Fig. 2 |
| SGR236 | *greEx217[sur-5p::HuSmad-4_NLS::MmPylRS::let-858_3’UTR; sur-5p::HygR::unc-54_3’UTR; rpr-1p::tRNA(M15)[UUA]::sup-7_3’UTR; rps-0p::GFP::TAG::mCherry::HA::EGL-13_NLS::unc-54_3’UTR]* | *smg-6(ok1794)* | JJ59(2), JJ268(7), SG88(3) | Fig. 3a |
| SGR237 | *greEx218[sur-5p::HuSmad-4_NLS::MmPylRS::let-858_3’UTR; sur-5p::HygR::unc-54_3’UTR; rpr-1p::tRNA(M15)[UCA]::sup-7_3’UTR; rps-0p::GFP::TAG::mCherry::HA::EGL-13_NLS::unc-54_3’UTR]* | *smg-6(ok1794)* | JJ59(2), KB337(7), SG88(3) | Fig. 3a |
| SGR238 | *greEx219[sur-5p::HuSmad-4_NLS::MmPylRS::let-858_3’UTR; sur-5p::HygR::unc-54_3’UTR; rpr-1p::tRNA(M15)::sup-7_3’UTR; rps-0p::GFP::TAA::mCherry::HA::EGL-13_NLS::unc-54_3’UTR]* | *smg-6(ok1794)* | JJ59(2), SE149(7), JJ266(3) | Fig. 3a |
| SGR239 | *greEx220[sur-5p::HuSmad-4_NLS::MmPylRS::let-858_3’UTR; sur-5p::HygR::unc-54_3’UTR; rpr-1p::tRNA(M15)[UCA]::sup-7_3’UTR; rps-0p::GFP::TAA::mCherry::HA::EGL-13_NLS::unc-54_3’UTR]* | *smg-6(ok1794)* | JJ59(2), KB337(7), JJ266(3) | Fig. 3a |
| SGR240 | *greEx221[sur-5p::HuSmad-4_NLS::MmPylRS::let-858_3’UTR; sur-5p::HygR::unc-54_3’UTR; rpr-1p::tRNA(M15)::sup-7_3’UTR; rps-0p::GFP::TGA::mCherry::HA::EGL-13_NLS::unc-54_3’UTR]* | *smg-6(ok1794)* | JJ59(2), SE149(7), JJ108(3) | Fig. 3a |
| SGR241 | *greEx222[sur-5p::HuSmad-4_NLS::MmPylRS::let-858_3’UTR; sur-5p::HygR::unc-54_3’UTR; rpr-1p::tRNA(M15)[UUA]::sup-7_3’UTR; rps-0p::GFP::TGA::mCherry::HA::EGL-13_NLS::unc-54_3’UTR]* | *smg-6(ok1794)* | JJ59(2), JJ268(7), JJ108(3) | Fig. 3a |
| SGR242 | *greEx223[sur-5p::HuSmad-4_NLS::MmPCKRS::let-858_3’UTR; rps-0p::HygR::unc-54_3’UTR; rpr-1p::tRNA(M15)::sup-7_3’UTR; rps-0p::GFP::TAG::mCherry::HA::EGL-13_NLS::unc-54_3’UTR]* | *smg-6(ok1794)* | SE170(2), SE149(6), SG88(3) | Fig. 3c |
| SGR243 | *greEx224[sur-5p::HuSmad-4_NLS::MmPCKRS::let-858_3’UTR; rps-0p::HygR::unc-54_3’UTR; rpr-1p::tRNA(M15)::sup-7_3’UTR; rps-0p::GFP::TAG::mCherry::HA::EGL-13_NLS::unc-54_3’UTR]* | *smg-6(ok1794)* | SE170(2), SE149(6), SG88(3) | Fig. 3c |
| SGR244 | *greEx225[sur-5p::HuSmad-4_NLS::MmPCKRS::let-858_3’UTR; rps-0p::HygR::unc-54_3’UTR; rpr-1p::tRNA(M15)[UACU,ev]::sup-7_3’UTR; rps-0p::GFP::AGTA::mCherry::HA::EGL-13_NLS::unc-54_3’UTR]* | *smg-6(ok1794)* | SE170(2), SE371(7), JJ458(3) | Fig. 3c |
| SGR245 | *greEx226[sur-5p::HuSmad-4_NLS::MmPCKRS::let-858_3’UTR; rps-0p::HygR::unc-54_3’UTR; rpr-1p::tRNA(M15)[UACU,ev]::sup-7_3’UTR; rps-0p::GFP::AGTA::mCherry::HA::EGL-13_NLS::unc-54_3’UTR]* | *smg-6(ok1794)* | SE170(2), SE371(7), JJ458(3) | Fig. 3c |
| SGR248 | *greEx229[sur-5p::HuSmad-4_NLS::MmPCKRS::let-858_3’UTR; rps-0p::HygR::unc-54_3’UTR; rpr-1p::tRNA(M15)[UCCU,M7]::sup-7_3’UTR; rps-0p::GFP::AGGA::mCherry::HA::EGL-13_NLS::unc-54_3’UTR]* | *smg-6(ok1794)* | SE170(2), SE289(7), JJ460(3) | Fig. 3c |
| SGR249 | *greEx230[sur-5p::HuSmad-4_NLS::MmPCKRS::let-858_3’UTR; rps-0p::HygR::unc-54_3’UTR; rpr-1p::tRNA(M15)[UCCU,M7]::sup-7_3’UTR; rps-0p::GFP::AGGA::mCherry::HA::EGL-13_NLS::unc-54_3’UTR]* | *smg-6(ok1794)* | SE170(2), SE289(7), JJ460(3) | Fig. 3c |
| SGR250 | *greEx231[sur-5p::HuSmad-4_NLS::MmPCKRS::let-858_3’UTR; rps-0p::HygR::unc-54_3’UTR; rpr-1p::tRNA(M15)[CUAG,ev]::sup-7_3’UTR; rps-0p::GFP::CTAG::mCherry::HA::EGL-13_NLS::unc-54_3’UTR]* | *smg-6(ok1794)* | SE170(2), SE370(7), JJ459(3) | Fig. 3c |
| SGR251 | *greEx232[sur-5p::HuSmad-4_NLS::MmPCKRS::let-858_3’UTR; rps-0p::HygR::unc-54_3’UTR; rpr-1p::tRNA(M15)[CUAG,ev]::sup-7_3’UTR; rps-0p::GFP::CTAG::mCherry::HA::EGL-13_NLS::unc-54_3’UTR]* | *smg-6(ok1794)* | SE170(2), SE370(7), JJ459(3) | Fig. 3c |
| SGR253 | *greEx234[sur-5p::MaPylRS::SL2::HuSmad-4_NLS::PCKRS::let-858_3’UTR; sur-5p::HygR::unc-54_3’UTR; rpr-1p::tRNA(M15)[CUA]::sup-7_3’UTR ; rpr-1p::tRNA(Ma10)[UCA]::sup-7_3’UTR; rps-0p::GFP::TGA::mCherry(D212TAG)::HA::EGL-13_NLS::unc-54_3’UTR]* | *smg-6(ok1794)* | JJ83(3), JJ357(7), JJ479(3) | Fig. 4b |
| SGR254 | *greEx235[sur-5p::MaPCKRS1::let-858_3’UTR; sur-5p::HygR::unc-54_3’UTR; rpr-1p::tRNA(Ma10)::sup-7_3’UTR; rps-0p::GFP::TAG::mCherry::HA::EGL-13_NLS::unc-54_3’UTR]* | *smg-6(ok1794)* | JJ85(2), SE298(7), SG88(3) | Fig. 5c |
| SGR255 | *greEx236[sur-5p::MaPCKRS1::let-858_3’UTR; sur-5p::HygR::unc-54_3’UTR; rpr-1p::tRNA(Ma10)::sup-7_3’UTR; rps-0p::GFP::TAG::mCherry::HA::EGL-13_NLS::unc-54_3’UTR]* | *smg-6(ok1794)* | JJ85(2), SE298(7), SG88(3) | Fig. 5c |
| SGR256 | *greEx237[sur-5p::MaPCKRS2::let-858_3’UTR; sur-5p::HygR::unc-54_3’UTR; rpr-1p::tRNA(Ma10)::sup-7_3’UTR; rps-0p::GFP::TAG::mCherry::HA::EGL-13_NLS::unc-54_3’UTR]* | *smg-6(ok1794)* | JJ186(2), SE298(7), SG88(3) | Fig. 5c |
| SGR257 | *greEx238[sur-5p::MaPCKRS2::let-858_3’UTR; sur-5p::HygR::unc-54_3’UTR; rpr-1p::tRNA(Ma10)::sup-7_3’UTR; rps-0p::GFP::TAG::mCherry::HA::EGL-13_NLS::unc-54_3’UTR]* | *smg-6(ok1794)* | JJ186(2), SE298(7), SG88(3) | Fig. 5c |
| SGR258 | *greEx239[sur-5p::MaPCKRS3::let-858_3’UTR; sur-5p::HygR::unc-54_3’UTR; rpr-1p::tRNA(Ma10)::sup-7_3’UTR; rps-0p::GFP::TAG::mCherry::HA::EGL-13_NLS::unc-54_3’UTR]* | *smg-6(ok1794)* | JJ243(2), SE298(7), SG88(3) | Fig. 5c |
| SGR259 | *greEx240[sur-5p::MaPCKRS3::let-858_3’UTR; sur-5p::HygR::unc-54_3’UTR; rpr-1p::tRNA(Ma10)::sup-7_3’UTR; rps-0p::GFP::TAG::mCherry::HA::EGL-13_NLS::unc-54_3’UTR]* | *smg-6(ok1794)* | JJ243(2), SE298(7), SG88(3) | Fig. 5c |
| SGR158 | *greEx143[sur-5p::HuSmad-4_NLS::HCKRS::let-858_3’UTR; sur-5p::HygR::unc-54_3’UTR; rpr-1p::tRNA(M15)::sup-7_3’UTR; rps-0p::GFP::TAG::mCherry::HA::EGL-13_NLS::unc-54_3’UTR]* | *smg-6(ok1794)* | JJ345(2), SE149(7), SG88(3) | Fig. 5c |
| SGR160 | *greEx145[sur-5p::HuSmad-4_NLS::HCKRS::let-858_3’UTR; sur-5p::HygR::unc-54_3’UTR; rpr-1p::tRNA(M15)::sup-7_3’UTR; rps-0p::GFP::TAG::mCherry::HA::EGL-13_NLS::unc-54_3’UTR]* | *smg-6(ok1794)* | JJ345(2), SE149(7), SG88(3) | Fig. 5c |
| SGR260 | *greEx241[sur-5p::MaHCKRS1::let-858_3’UTR; sur-5p::HygR::unc-54_3’UTR; rpr-1p::tRNA(Ma10)::sup-7_3’UTR; rps-0p::GFP::TAG::mCherry::HA::EGL-13_NLS::unc-54_3’UTR]* | *smg-6(ok1794)* | JJ347(2), SE298(7), SG88(3) | Fig. 5c |
| SGR261 | *greEx242[sur-5p::MaHCKRS2::let-858_3’UTR; sur-5p::HygR::unc-54_3’UTR; rpr-1p::tRNA(Ma10)::sup-7_3’UTR; rps-0p::GFP::TAG::mCherry::HA::EGL-13_NLS::unc-54_3’UTR]* | *smg-6(ok1794)* | JJ348(2), SE298(7), SG88(3) | Fig. 5c |
| SGR262 | *greEx243[sur-5p::MaHCKRS2::let-858_3’UTR; sur-5p::HygR::unc-54_3’UTR; rpr-1p::tRNA(Ma10)::sup-7_3’UTR; rps-0p::GFP::TAG::mCherry::HA::EGL-13_NLS::unc-54_3’UTR]* | *smg-6(ok1794)* | JJ348(2), SE298(7), SG88(3) | Fig. 5c |
| SGR263 | *greEx244[sur-5p::MaHCKRS3::let-858_3’UTR; sur-5p::HygR::unc-54_3’UTR; rpr-1p::tRNA(Ma10)::sup-7_3’UTR; rps-0p::GFP::TAG::mCherry::HA::EGL-13_NLS::unc-54_3’UTR]* | *smg-6(ok1794)* | JJ349(2), SE298(7), SG88(3) | Fig. 5c |
| SGR264 | *greEx245[sur-5p::MaHCKRS3::let-858_3’UTR; sur-5p::HygR::unc-54_3’UTR; rpr-1p::tRNA(Ma10)::sup-7_3’UTR; rps-0p::GFP::TAG::mCherry::HA::EGL-13_NLS::unc-54_3’UTR]* | *smg-6(ok1794)* | JJ349(2), SE298(7), SG88(3) | Fig. 5c |
| SGR265 | *greEx246[sur-5p::MaHCKRS3::let-858_3’UTR; sur-5p::HygR::unc-54_3’UTR; rpr-1p::tRNA(Ma10)::sup-7_3’UTR; rps-0p::GFP::TAG::mCherry::HA::EGL-13_NLS::unc-54_3’UTR]* | *smg-6(ok1794)* | JJ349(2), SE298(7), SG88(3) | Fig. 5c |
| SGR266 | *greEx247[sur-5p::HuSmad-4_NLS::NPYRS::let-858_3’UTR; rps-0p::HygR::unc-54_3’UTR; rpr-1p::tRNA(M15)::sup-7_3’UTR; rps-0p::GFP::TAG::mCherry::HA::EGL-13_NLS::unc-54_3’UTR]* | *smg-6(ok1794)* | JOS98(2), SE149(7), SG88(3) | Fig. 5b |
| SGR267 | *greEx248[sur-5p::HuSmad-4_NLS::NPYRS::let-858_3’UTR; rps-0p::HygR::unc-54_3’UTR; rpr-1p::tRNA(M15)::sup-7_3’UTR; rps-0p::GFP::TAG::mCherry::HA::EGL-13_NLS::unc-54_3’UTR]* | *smg-6(ok1794)* | JOS98(2), SE149(7), SG88(3) | Fig. 5b |
| SGR268 | *greEx249[sur-5p::MaNPYRS1::let-858_3’UTR; sur-5p::HygR::unc-54_3’UTR; rpr-1p::tRNA(Ma10)::sup-7_3’UTR; rps-0p::GFP::TAG::mCherry::HA::EGL-13_NLS::unc-54_3’UTR]* | *smg-6(ok1794)* | JJ245(2), SE298(7), SG88(3) | Fig. 5b |
| SGR269 | *greEx250[sur-5p::MaNPYRS1::let-858_3’UTR; sur-5p::HygR::unc-54_3’UTR; rpr-1p::tRNA(Ma10)::sup-7_3’UTR; rps-0p::GFP::TAG::mCherry::HA::EGL-13_NLS::unc-54_3’UTR]* | *smg-6(ok1794)* | JJ245(2), SE298(7), SG88(3) | Fig. 5b |
| SGR270 | *greEx251[sur-5p::MaNPYRS2::let-858_3’UTR; sur-5p::HygR::unc-54_3’UTR; rpr-1p::tRNA(Ma10)::sup-7_3’UTR; rps-0p::GFP::TAG::mCherry::HA::EGL-13_NLS::unc-54_3’UTR]* | *smg-6(ok1794)* | JJ189(2), SE298(7), SG88(3) | Fig. 5b |
| SGR271 | *greEx252[sur-5p::MaNPYRS2::let-858_3’UTR; sur-5p::HygR::unc-54_3’UTR; rpr-1p::tRNA(Ma10)::sup-7_3’UTR; rps-0p::GFP::TAG::mCherry::HA::EGL-13_NLS::unc-54_3’UTR]* | *smg-6(ok1794)* | JJ189(2), SE298(7), SG88(3) | Fig. 5b |
| SGR272 | *greEx253[sur-5p::MaNPYRS3::let-858_3’UTR; sur-5p::HygR::unc-54_3’UTR; rpr-1p::tRNA(Ma10)::sup-7_3’UTR; rps-0p::GFP::TAG::mCherry::HA::EGL-13_NLS::unc-54_3’UTR]* | *smg-6(ok1794)* | JJ241(2), SE298(7), SG88(3) | Fig. 5b |
| SGR273 | *greEx254[sur-5p::MaNPYRS3::let-858_3’UTR; sur-5p::HygR::unc-54_3’UTR; rpr-1p::tRNA(Ma10)::sup-7_3’UTR; rps-0p::GFP::TAG::mCherry::HA::EGL-13_NLS::unc-54_3’UTR]* | *smg-6(ok1794)* | JJ241(2), SE298(7), SG88(3) | Fig. 5b |
| SGR274 | *greEx255[sur-5p::HuSmad-4_NLS::PCC2RS::let-858_3’UTR; sur-5p::HygR::unc-54_3’UTR; rpr-1p::tRNA(M15)::sup-7_3’UTR; rps-0p::GFP::TAG::mCherry::HA::EGL-13_NLS::unc-54_3’UTR]* | *smg-6(ok1794)* | JJ579(2), SE149(7), SG88(3) | Fig. 5d |
| SGR275 | *greEx256[sur-5p::HuSmad-4_NLS::PCC2RS::let-858_3’UTR; sur-5p::HygR::unc-54_3’UTR; rpr-1p::tRNA(M15)::sup-7_3’UTR; rps-0p::GFP::TAG::mCherry::HA::EGL-13_NLS::unc-54_3’UTR]* | *smg-6(ok1794)* | JJ579(2), SE149(7), SG88(3) | Fig. 5d |
| SGR276 | *greEx257[sur-5p::MaPCC2RS1::let-858_3’UTR; sur-5p::HygR::unc-54_3’UTR; rpr-1p::tRNA(Ma10)::sup-7_3’UTR; rps-0p::GFP::TAG::mCherry::HA::EGL-13_NLS::unc-54_3’UTR]* | *smg-6(ok1794)* | JJ550(2), SE298(7), SG88(3) | Fig. 5d |
| SGR277 | *greEx258[sur-5p::MaPCC2RS1::let-858_3’UTR; sur-5p::HygR::unc-54_3’UTR; rpr-1p::tRNA(Ma10)::sup-7_3’UTR; rps-0p::GFP::TAG::mCherry::HA::EGL-13_NLS::unc-54_3’UTR]* | *smg-6(ok1794)* | JJ550(2), SE298(7), SG88(3) | Fig. 5d |
| SGR278 | *greEx259[sur-5p::MaPCC2RS2::let-858_3’UTR; sur-5p::HygR::unc-54_3’UTR; rpr-1p::tRNA(Ma10)::sup-7_3’UTR; rps-0p::GFP::TAG::mCherry::HA::EGL-13_NLS::unc-54_3’UTR]* | *smg-6(ok1794)* | JJ551(2), SE298(7), SG88(3) | Fig. 5d |
| SGR279 | *greEx260[sur-5p::MaPCC2RS2::let-858_3’UTR; sur-5p::HygR::unc-54_3’UTR; rpr-1p::tRNA(Ma10)::sup-7_3’UTR; rps-0p::GFP::TAG::mCherry::HA::EGL-13_NLS::unc-54_3’UTR]* | *smg-6(ok1794)* | JJ551(2), SE298(7), SG88(3) | Fig. 5d |
| SGR280 | *greEx261[sur-5p::MaPCC2RS2::let-858_3’UTR; sur-5p::HygR::unc-54_3’UTR; rpr-1p::tRNA(Ma10)::sup-7_3’UTR; rps-0p::GFP::TAG::mCherry::HA::EGL-13_NLS::unc-54_3’UTR]* | *smg-6(ok1794)* | JJ551(2), SE298(7), SG88(3) | Fig. 5d |
| SGR281 | *greEx262[sur-5p::MaPCC2RS3::let-858_3’UTR; sur-5p::HygR::unc-54_3’UTR; rpr-1p::tRNA(Ma10)::sup-7_3’UTR; rps-0p::GFP::TAG::mCherry::HA::EGL-13_NLS::unc-54_3’UTR]* | *smg-6(ok1794)* | JJ552(2), SE298(7), SG88(3) | Fig. 5d |
| SGR282 | *greEx263[sur-5p::MaPCC2RS3::let-858_3’UTR; sur-5p::HygR::unc-54_3’UTR; rpr-1p::tRNA(Ma10)::sup-7_3’UTR; rps-0p::GFP::TAG::mCherry::HA::EGL-13_NLS::unc-54_3’UTR]* | *smg-6(ok1794)* | JJ552(2), SE298(7), SG88(3) | Fig. 5d |
| SGR283 | *greEx264[sur-5p::MaPCC2RS3::let-858_3’UTR; sur-5p::HygR::unc-54_3’UTR; rpr-1p::tRNA(Ma10)::sup-7_3’UTR; rps-0p::GFP::TAG::mCherry::HA::EGL-13_NLS::unc-54_3’UTR]* | *smg-6(ok1794)* | JJ552(2), SE298(7), SG88(3) | Fig. 5d |
| SGR284 | *greEx265[sur-5p::MaPCC2RS3::let-858_3’UTR; sur-5p::HygR::unc-54_3’UTR; rpr-1p::tRNA(Ma10)::sup-7_3’UTR; rps-0p::GFP::TAG::mCherry::HA::EGL-13_NLS::unc-54_3’UTR]* | *smg-6(ok1794)* | JJ552(2), SE298(7), SG88(3) | Fig. 5d |
| SGR141 | *greEx126[glr-1p::HuSmad-4_NLS::NPYRS::let-858_3’UTR; sur-5p::HygR::unc-54_3’UTR; rpr-1p::tRNA(M15)::sup-7_3’UTR; glr-1p::Cre(R119A,Y324TAG)::HA::EGL-13_NLS::let-858_3’UTR; maco-1p::loxP::B-Gal_3'UTR::loxP::Citrine2::HA::let-858_3’UTR]* | *N2* | AT11(1.5), SE149(4.5), JJ191(6), KB462(2.5) | Fig. 6d, Fig. 7a |
| SGR142 | *greEx127[glr-1p::HuSmad-4_NLS::NPYRS::let-858_3’UTR; sur-5p::HygR::unc-54_3’UTR; rpr-1p::tRNA(M15)::sup-7_3’UTR; glr-1p::Cre(R119A,Y324TAG)::HA::EGL-13_NLS::let-858_3’UTR; maco-1p::loxP::B-Gal_3'UTR::loxP::Citrine2::HA::let-858_3’UTR]* | *N2* | AT11(1.5), SE149(4.5), JJ191(6), KB462(2.5) | Fig. 6d, Fig. 7a |
| SGR285 | *greEx266[glr-1p::MaNPYRS2::let-858_3’UTR; sur-5p::HygR::unc-54_3’UTR; rpr-1p::tRNA(Ma10)::sup-7_3’UTR; glr-1p::FLP(Y343TAG)::HA::EGL-13_NLS::SL2::mKate2::let-858_3’UTR; maco-1p::FRT::B-Gal_3'UTR::FRT::Citrine2::HA::let-858_3’UTR]* | *N2* | JJ244(1.5), SE298(4.5), KB463(6), KB454(2.5) | Fig. 6c, d |
| SGR286 | *greEx267[glr-1p::MaNPYRS2::let-858_3’UTR; sur-5p::HygR::unc-54_3’UTR; rpr-1p::tRNA(Ma10)::sup-7_3’UTR; glr-1p::FLP(Y343TAG)::HA::EGL-13_NLS::SL2::mKate2::let-858_3’UTR; maco-1p::FRT::B-Gal_3'UTR::FRT::Citrine2::HA::let-858_3’UTR]* | *N2* | JJ244(1.5), SE298(4.5), KB463(6), KB454(2.5) | Fig. 6d |
| SGR287 | *greEx268[glr-1p::MaNPYRS2::let-858_3’UTR; sur-5p::HygR::unc-54_3’UTR; rpr-1p::tRNA(Ma10)::sup-7_3’UTR; glr-1p::FLP(Y343TAG)::HA::EGL-13_NLS::SL2::mKate2::let-858_3’UTR; maco-1p::FRT::B-Gal_3'UTR::FRT::Citrine2::HA::let-858_3’UTR]* | *N2* | JJ244(1.5), SE298(4.5), KB463(6), KB454(2.5) | Fig. 6d |
| SGR288 | *greEx269[glr-1p::MaNPYRS2::let-858_3’UTR; sur-5p::HygR::unc-54_3’UTR; rpr-1p::tRNA(Ma10)::sup-7_3’UTR; glr-1p::Cre(R119A,Y324TAG)::HA::EGL-13_NLS::let-858_3’UTR; maco-1p::FRT::B-Gal_3'UTR::FRT::Citrine2::HA::let-858_3’UTR]* | *N2* | JJ244(1.5), SE298(4.5), JJ191(6), KB462(2.5) | Fig. 6d |
| SGR289 | *greEx270[glr-1p::MaNPYRS2::let-858_3’UTR; sur-5p::HygR::unc-54_3’UTR; rpr-1p::tRNA(Ma10)::sup-7_3’UTR; glr-1p::Cre(R119A,Y324TAG)::HA::EGL-13_NLS::let-858_3’UTR; maco-1p::FRT::B-Gal_3'UTR::FRT::Citrine2::HA::let-858_3’UTR]* | *N2* | JJ244(1.5), SE298(4.5), JJ191(6), KB462(2.5) | Fig. 6d |
| SGR290 | *greEx271[glr-1p::MaNPYRS2::let-858_3’UTR; sur-5p::HygR::unc-54_3’UTR; rpr-1p::tRNA(Ma10)::sup-7_3’UTR; glr-1p::Cre(R119A,Y324TAG)::HA::EGL-13_NLS::let-858_3’UTR; maco-1p::FRT::B-Gal_3'UTR::FRT::Citrine2::HA::let-858_3’UTR]* | *N2* | JJ244(1.5), SE298(4.5), JJ191(6), KB462(2.5) | Fig. 6d |
| SGR291 | *greEx272[glr-1p::MaNPYRS3::let-858_3’UTR; sur-5p::HygR::unc-54_3’UTR; rpr-1p::tRNA(Ma10)::sup-7_3’UTR; glr-1p::FLP(Y343TAG)::HA::EGL-13_NLS::SL2::mKate2::let-858_3’UTR; maco-1p::FRT::B-Gal_3'UTR::FRT::Citrine2::HA::let-858_3’UTR]* | *N2* | JJ446(1.5), SE298(4.5), KB463(6), KB454(2.5) | Fig. 6c, d |
| SGR292 | *greEx273[glr-1p::MaNPYRS3::let-858_3’UTR; sur-5p::HygR::unc-54_3’UTR; rpr-1p::tRNA(Ma10)::sup-7_3’UTR; glr-1p::FLP(Y343TAG)::HA::EGL-13_NLS::SL2::mKate2::let-858_3’UTR; maco-1p::FRT::B-Gal_3'UTR::FRT::Citrine2::HA::let-858_3’UTR]* | *N2* | JJ446(1.5), SE298(4.5), KB463(6), KB454(2.5) | Fig. 6d |
| SGR293 | *greEx274[glr-1p::MaNPYRS3::let-858_3’UTR; sur-5p::HygR::unc-54_3’UTR; rpr-1p::tRNA(Ma10)::sup-7_3’UTR; glr-1p::Cre(R119A,Y324TAG)::HA::EGL-13_NLS::let-858_3’UTR; maco-1p::loxP::B-Gal_3'UTR::loxP::Citrine2::HA::let-858_3’UTR]* | *N2* | JJ446(1.5), SE298(4.5), JJ191(6), KB462(2.5) | Fig. 6d |
| SGR294 | *greEx275[glr-1p::MaNPYRS3::let-858_3’UTR; sur-5p::HygR::unc-54_3’UTR; rpr-1p::tRNA(Ma10)::sup-7_3’UTR; glr-1p::Cre(R119A,Y324TAG)::HA::EGL-13_NLS::let-858_3’UTR; maco-1p::loxP::B-Gal_3'UTR::loxP::Citrine2::HA::let-858_3’UTR]* | *N2* | JJ446(1.5), SE298(4.5), JJ191(6), KB462(2.5) | Fig. 6d |
| SGR295 | *greEx276[glr-1p::MaPCC2RS1::let-858_3’UTR; sur-5p::HygR::unc-54_3’UTR; rpr-1p::tRNA(Ma10)::sup-7_3’UTR; glr-1p::FLP(C189TAG)::HA::EGL-13_NLS::SL2::mKate2::let-858_3’UTR; maco-1p::FRT::B-Gal_3'UTR::FRT::Citrine2::HA::let-858_3’UTR]* | *N2* | JJ629(1.5), SE298(4.5), JJ543(6), KB454(2.5) | Fig. 6c, e, Fig. 7a |
| SGR296 | *greEx277[glr-1p::MaPCC2RS1::let-858_3’UTR; sur-5p::HygR::unc-54_3’UTR; rpr-1p::tRNA(Ma10)::sup-7_3’UTR; glr-1p::FLP(C189TAG)::HA::EGL-13_NLS::SL2::mKate2::let-858_3’UTR; maco-1p::FRT::B-Gal_3'UTR::FRT::Citrine2::HA::let-858_3’UTR]* | *N2* | JJ629(1.5), SE298(4.5), JJ543(6), KB454(2.5) | Fig. 6e, Fig. 7a |
| SGR297 | *greEx278[glr-1p::MaPCC2RS1::let-858_3’UTR; sur-5p::HygR::unc-54_3’UTR; rpr-1p::tRNA(Ma10)::sup-7_3’UTR; glr-1p::FLP(C189TAG)::HA::EGL-13_NLS::SL2::mKate2::let-858_3’UTR; maco-1p::FRT::B-Gal_3'UTR::FRT::Citrine2::HA::let-858_3’UTR]* | *N2* | JJ629(1.5), SE298(4.5), JJ543(6), KB454(2.5) | Fig. 6e, Fig. 7a |
| SGR298 | *greEx279[glr-1p::MaPCC2RS1::let-858_3’UTR; sur-5p::HygR::unc-54_3’UTR; rpr-1p::tRNA(Ma10)::sup-7_3’UTR; glr-1p::FLP(C189TAG)::HA::EGL-13_NLS::SL2::mKate2::let-858_3’UTR; maco-1p::FRT::B-Gal_3'UTR::FRT::Citrine2::HA::let-858_3’UTR]* | *N2* | JJ629(1.5), SE298(4.5), JJ543(6), KB454(2.5) | Fig. 6e, Fig. 7a |
| SGR299 | *greEx280[glr-1p::MaPCC2RS1::let-858_3’UTR; sur-5p::HygR::unc-54_3’UTR; rpr-1p::tRNA(Ma10)::sup-7_3’UTR; glr-1p::FLP(C189TAG)::HA::EGL-13_NLS::SL2::mKate2::let-858_3’UTR; maco-1p::FRT::B-Gal_3'UTR::FRT::Citrine2::HA::let-858_3’UTR]* | *N2* | JJ629(1.5), SE298(4.5), JJ543(6), KB454(2.5) | Fig. 6e, Fig. 7a |
| SGR300 | *greEx281[glr-1p::MaPCC2RS2::let-858_3’UTR; sur-5p::HygR::unc-54_3’UTR; rpr-1p::tRNA(Ma10)::sup-7_3’UTR; glr-1p::FLP(C189TAG)::HA::EGL-13_NLS::SL2::mKate2::let-858_3’UTR; maco-1p::FRT::B-Gal_3'UTR::FRT::Citrine2::HA::let-858_3’UTR]* | *N2* | JJ575(1.5), SE298(4.5), JJ543(6), KB454(2.5) | Fig. 6e |
| SGR301 | *greEx282[glr-1p::MaPCC2RS2::let-858_3’UTR; sur-5p::HygR::unc-54_3’UTR; rpr-1p::tRNA(Ma10)::sup-7_3’UTR; glr-1p::FLP(C189TAG)::HA::EGL-13_NLS::SL2::mKate2::let-858_3’UTR; maco-1p::FRT::B-Gal_3'UTR::FRT::Citrine2::HA::let-858_3’UTR]* | *N2* | JJ575(1.5), SE298(4.5), JJ543(6), KB454(2.5) | Fig. 6e |
| SGR302 | *greEx283[glr-1p::MaPCC2RS2::let-858_3’UTR; sur-5p::HygR::unc-54_3’UTR; rpr-1p::tRNA(Ma10)::sup-7_3’UTR; glr-1p::FLP(C189TAG)::HA::EGL-13_NLS::SL2::mKate2::let-858_3’UTR; maco-1p::FRT::B-Gal_3'UTR::FRT::Citrine2::HA::let-858_3’UTR]* | *N2* | JJ575(1.5), SE298(4.5), JJ543(6), KB454(2.5) | Fig. 6e |
| SGR303 | *greEx284[glr-1p::MaPCC2RS3::let-858_3’UTR; sur-5p::HygR::unc-54_3’UTR; rpr-1p::tRNA(Ma10)::sup-7_3’UTR; glr-1p::FLP(C189TAG)::HA::EGL-13_NLS::SL2::mKate2::let-858_3’UTR; maco-1p::FRT::B-Gal_3'UTR::FRT::Citrine2::HA::let-858_3’UTR]* | *N2* | JJ630(1.5), SE298(4.5), JJ543(6), KB454(2.5) | Fig. 6e |
| SGR304 | *greEx285[glr-1p::MaPCC2RS3::let-858_3’UTR; sur-5p::HygR::unc-54_3’UTR; rpr-1p::tRNA(Ma10)::sup-7_3’UTR; glr-1p::FLP(C189TAG)::HA::EGL-13_NLS::SL2::mKate2::let-858_3’UTR; maco-1p::FRT::B-Gal_3'UTR::FRT::Citrine2::HA::let-858_3’UTR]* | *N2* | JJ630(1.5), SE298(4.5), JJ543(6), KB454(2.5) | Fig. 6e |
| SGR305 | *greEx286[glr-1p::MaPCC2RS3::let-858_3’UTR; sur-5p::HygR::unc-54_3’UTR; rpr-1p::tRNA(Ma10)::sup-7_3’UTR; glr-1p::FLP(C189TAG)::HA::EGL-13_NLS::SL2::mKate2::let-858_3’UTR; maco-1p::FRT::B-Gal_3'UTR::FRT::Citrine2::HA::let-858_3’UTR]* | *N2* | JJ630(1.5), SE298(4.5), JJ543(6), KB454(2.5) | Fig. 6e |
| SGR306 | *greEx287[glr-1p::MaPCC2RS3::let-858_3’UTR; sur-5p::HygR::unc-54_3’UTR; rpr-1p::tRNA(Ma10)::sup-7_3’UTR; glr-1p::FLP(C189TAG)::HA::EGL-13_NLS::SL2::mKate2::let-858_3’UTR; maco-1p::FRT::B-Gal_3'UTR::FRT::Citrine2::HA::let-858_3’UTR]* | *N2* | JJ630(1.5), SE298(4.5), JJ543(6), KB454(2.5) | Fig. 6e |
| SGR307 | *greEx288[glr-1p::HuSmad-4_NLS::NPYRS::SL2::4x(rpr-1p::tRNA(M15)::sup-7_3’UTR)::let-858_3’UTR; sur-5p::HygR::unc-54_3’UTR; glr-1p::MaPCC2RS1::SL2::4x(rpr-1p::tRNA(Ma10)[UCA]::sup-7_3’UTR)::let-858_3’UTR; sur-5p::HygR::unc-54_3’UTR; glr-1p::FLP(C189TGA)::HA::EGL-13_NLS::SL2::Cre(R119A,Y324TAG)::HA::EGL-13_NLS::SL2::CFP::let-858_3’UTR; maco-1p::FRT::B-Gal_3'UTR::FRT::Citrine2::HA::let-858_3’UTR; maco-1p::loxP::B-Gal_3'UTR::loxP::mKate2::let-858_3’UTR]* | *N2* | JJ676(3.5), JJ681(3.5), JJ685(7), KB454(2.5), JJ713(2.5) | Fig. 7d |
| SGR308 | *greEx289[glr-1p::HuSmad-4_NLS::NPYRS::SL2::4x(rpr-1p::tRNA(M15)::sup-7_3’UTR)::let-858_3’UTR; sur-5p::HygR::unc-54_3’UTR; glr-1p::MaPCC2RS1::SL2::4x(rpr-1p::tRNA(Ma10)[UCA]::sup-7_3’UTR)::let-858_3’UTR; sur-5p::HygR::unc-54_3’UTR; glr-1p::FLP(C189TGA)::HA::EGL-13_NLS::SL2::Cre(R119A,Y324TAG)::HA::EGL-13_NLS::SL2::CFP::let-858_3’UTR; maco-1p::FRT::B-Gal_3'UTR::FRT::Citrine2::HA::let-858_3’UTR; maco-1p::loxP::B-Gal_3'UTR::loxP::mKate2::let-858_3’UTR]* | *N2* | JJ676(3.5), JJ681(3.5), JJ685(7), KB454(2.5), JJ713(2.5) | Fig. 7d |

**Supplementary Table 2 Expression plasmids.**

| Plasmid  Number | Description | Source Plasmids | | | |
| --- | --- | --- | --- | --- | --- |
|  |  | **Dest** | **p1** | **p2** | **p3** |
| AT11 | *glr-1p::HuSmad-4_NLS::NPYRS::let-858_3’UTR; sur-5p::HygR::unc-54_3’UTR* | SE80 | IR157 | KB126 | SG606 |
| JJ57 | *glr-1p::MaPCC2RS2::let-858_3’UTR; sur-5p::HygR::unc-54_3’UTR* | SE80 | IR157 | JJ548 | SG606 |
| JJ59 | *sur-5p::HuSmad-4_NLS::MmPylRS::let-858_3’UTR; sur-5p::HygR::unc-54_3’UTR* | SE80 | SE72 | JJ58 | SG606 |
| JJ60 | *sur-5p::FLAG::G1PylRS::let-858_3’UTR; sur-5p::HygR::unc-54_3’UTR* | SE80 | SE72 | JJ56 | SG606 |
| JJ83 | sur-5p::MaPylRS::SL2::HuSmad-4_NLS::PCKRS::let-858_3’UTR; sur-5p::HygR::unc-54_3’UTR | SE80 | SE72 | KB182 | JJ82 |
| JJ85 | *sur-5p::MaPCKRS1::let-858_3’UTR; sur-5p::HygR::unc-54_3’UTR* | SE80 | SE72 | JJ84 | SG606 |
| JJ108 | *rps-0p::GFP::TGA::mCherry::HA::EGL-13_NLS::unc-54_3’UTR* | N/A | | | |
| JJ186 | *sur-5p::MaPCKRS2::let-858_3’UTR; sur-5p::HygR::unc-54_3’UTR* | SE80 | SE72 | JJ185 | SG606 |
| JJ189 | *sur-5p::MaNPYRS2::let-858_3’UTR; sur-5p::HygR::unc-54_3’UTR* | SE80 | SE72 | JJ188 | SG606 |
| JJ191 | *glr-1p::Cre(R119A,Y324TAG)::HA::EGL-13_NLS::let-858_3’UTR* | pDEST R4-R3 Vector II | IR157 | JJ190 | SG606 |
| JJ241 | *sur-5p::MaNPYRS3::let-858_3’UTR; sur-5p::HygR::unc-54_3’UTR* | SE80 | SE72 | JJ240 | SG606 |
| JJ243 | *sur-5p::MaPCKRS3::let-858_3’UTR; sur-5p::HygR::unc-54_3’UTR* | SE80 | SE72 | JJ242 | SG606 |
| JJ244 | *glr-1p::MaNPYRS2::let-858_3’UTR; sur-5p::HygR::unc-54_3’UTR* | SE80 | IR157 | JJ188 | SG606 |
| JJ245 | *sur-5p::MaNPYRS1::let-858_3’UTR; sur-5p::HygR::unc-54_3’UTR* | SE80 | SE72 | JJ239 | SG606 |
| JJ266 | *rps-0p::GFP::TAA::mCherry::HA::EGL-13_NLS::unc-54_3’UTR* | N/A | | | |
| JJ345 | *sur-5p::HuSmad-4_NLS::HCKRS::let-858_3’UTR; sur-5p::HygR::unc-54_3’UTR* | SE80 | SE72 | JJ338 | SG606 |
| JJ347 | *sur-5p::MaHCKRS1::let-858_3’UTR; sur-5p::HygR::unc-54_3’UTR* | SE80 | SE72 | JJ340 | SG606 |
| JJ348 | *sur-5p::MaHCKRS2::let-858_3’UTR; sur-5p::HygR::unc-54_3’UTR* | SE80 | SE72 | JJ341 | SG606 |
| JJ349 | *sur-5p::MaHCKRS3::let-858_3’UTR; sur-5p::HygR::unc-54_3’UTR* | SE80 | SE72 | JJ342 | SG606 |
| JJ446 | *glr-1p::MaNPYRS3::let-858_3’UTR; sur-5p::HygR::unc-54_3’UTR* | SE80 | IR157 | JJ240 | SG606 |
| JJ466 | *glr-1p::HuSmad-4_NLS::PCKRS(13IPYER)::let-858_3’UTR; sur-5p::HygR::unc-54_3’UTR* | SE80 | IR157 | JJ237 | SG606 |
| JJ518 | *sur-5p::HuSmad-4_NLS::MmPCKRS::let-858_3’UTR; sur-5p::HygR::unc-54_3’UTR* | SE80 | SE72 | SE154 | SG606 |
| JJ543 | *glr-1p::FLP(C189TAG)::HA::EGL-13_NLS::SL2::mKate2::let-858_3’UTR* | pDEST R4-R3 Vector II | IR157 | JJ540 | IR322 |
| JJ546 | *glr-1p::HuSmad-4_NLS::PCC2RS::let-858_3’UTR; sur-5p::HygR::unc-54_3’UTR* | SE80 | IR157 | KB124 | SG606 |
| JJ550 | *sur-5p::MaPCC2RS1::let-858_3’UTR; sur-5p::HygR::unc-54_3’UTR* | SE80 | SE72 | JJ547 | SG606 |
| JJ551 | *sur-5p::MaPCC2RS2::let-858_3’UTR; sur-5p::HygR::unc-54_3’UTR* | SE80 | SE72 | JJ548 | SG606 |
| JJ552 | *sur-5p::MaPCC2RS3::let-858_3’UTR; sur-5p::HygR::unc-54_3’UTR* | SE80 | SE72 | JJ549 | SG606 |
| JJ575 | *glr-1p::MaPCC2RS2::let-858_3’UTR; sur-5p::HygR::unc-54_3’UTR* | SE80 | IR157 | JJ548 | SG606 |
| JJ579 | *sur-5p::HuSmad-4_NLS::PCC2RS::let-858_3’UTR; sur-5p::HygR::unc-54_3’UTR* | SE80 | SE72 | KB124 | SG606 |
| JJ629 | *glr-1p::MaPCC2RS1::let-858_3’UTR; sur-5p::HygR::unc-54_3’UTR* | SE80 | IR157 | JJ547 | SG606 |
| JJ630 | *glr-1p::MaPCC2RS3::let-858_3’UTR; sur-5p::HygR::unc-54_3’UTR* | SE80 | IR157 | JJ549 | SG606 |
| JJ676 | *glr-1p::HuSmad-4_NLS::NPYRS::SL2::4x(rpr-1p::tRNA(M15)::sup-7_3’UTR)::let-858_3’UTR; sur-5p::HygR::unc-54_3’UTR* | SE80 | IR157 | KB126 | JJ472 |
| JJ681 | *glr-1p::MaPCC2RS1::SL2::4x(rpr-1p::tRNA(Ma10)[UCA]::sup-7_3’UTR)::let-858_3’UTR; sur-5p::HygR::unc-54_3’UTR* | SE80 | IR157 | JJ547 | JJ680 |
| JJ685 | *glr-1p::FLP(C189TGA)::HA::EGL-13_NLS::SL2::Cre(R119A,Y324TAG)::HA::EGL-13_NLS::SL2::CFP::let-858_3’UTR* | JJ331 | IR157 | JJ683 | JJ271 |
| JJ713 | *maco-1p::loxP::lacZ:loxP::mKate2::let-858_3’UTR* | pDEST R4-R3 Vector II | SE77 | SG365 | JJ426 |
| JOS98 | *sur-5p::HuSmad-4_NLS::NPYRS::let-858_3’UTR; rps-0p::HygR::unc-54_3’UTR* | IR98 | SE72 | KB126 | SG606 |
| KB375 | *sur-5p::MaPylRS::let-858_3’UTR; sur-5p::HygR::unc-54_3’UTR* | SE80 | SE72 | KB182 | SG606 |
| KB438 | *glr-1p::FLP(K223TAG)::HA::EGL-13_NLS::SL2::mKate2::let-858_3’UTR* | pDEST R4-R3 Vector II | IR157 | KB257 | IR322 |
| KB454 | *maco-1p::FRT::lacZ::FRT::Citrine2::HA::let-858_3’UTR* | pDEST R4-R3 Vector II | SE77 | LD222 | KB442 |
| KB462 | *maco-1p::loxP::lacZ:loxP::Citrine2::HA::let-858_3’UTR* | pDEST R4-R3 Vector II | SE77 | SG365 | KB442 |
| KB463 | *glr-1p::FLP(Y343TAG)::HA::EGL-13_NLS::SL2::mKate2::let-858_3’UTR* | pDEST R4-R3 Vector II | IR157 | KB321 | IR322 |
| SG88 | *rps-0p::GFP::TAG::mCherry::HA::EGL-13_NLS::unc-54_3’UTR* | ^1^ | | | |
| JJ267 | *rpr-1p::tRNA(Ma10)[UUA]::sup-7_3’UTR* | N/A | | | |
| JJ268 | *rpr-1p::tRNA(M15)[UUA]::sup-7_3’UTR* | N/A | | | |
| JJ269 | *rpr-1p::tRNA(Ma10)[UCA]::sup-7_3’UTR* | N/A | | | |
| KB337 | *rpr-1p::tRNA(M15)[UCA]::sup-7_3’UTR* | N/A | | | |
| SE149 | *rpr-1p::tRNA(M15)::sup-7_3’UTR* | ^2^ | | | |
| SE150 | *rpr-1p::tRNA(C15)::sup-7_3’UTR* | ^3^ | | | |
| SE297 | *rpr-1p::tRNA(Ma6)::sup-7_3’UTR* | N/A | | | |
| SE298 | *rpr-1p::tRNA(Ma10)::sup-7_3’UTR* | N/A | | | |

**Supplementary Table 3 Entry plasmids.**

| Plasmid Number | Description | Source |
| --- | --- | --- |
| IR98 | *rps-0p::HygR::unc-54_3’UTR* | ^3^ |
| IR157 | *glr-1p* | ^3^ |
| IR322 | *SL2::mKate2::let-858_3’UTR* | ^4^ |
| JJ56 | *FLAG::G1PylRS* | This paper |
| JJ58 | *HuSmad-4_NLS::MmPylRS* | This paper |
| JJ84 | *MaPCKRS1* | This paper |
| JJ185 | *MaPCKRS2* | This paper |
| JJ188 | *MaNPYRS2* | This paper |
| JJ190 | *Cre(R119A,Y324TAG)::HA::EGL-13_NLS* | ^4^ |
| JJ237 | *HuSmad-4_NLS::PCKRS(13IPYER)* | ^4^ |
| JJ239 | *MaNPYRS1* | This paper |
| JJ240 | *MaNPYRS3* | This paper |
| JJ242 | *MaPCKRS3* | This paper |
| JJ271 | *SL2::Cre(R119A,Y324TAG)::HA::EGL-13_NLS* | This paper |
| JJ331 | *SL2::CFP::let-858_3’UTR* | This paper |
| JJ338 | *HuSmad-4_NLS::HCKRS* | ^4^ |
| JJ340 | *MaHCKRS1* | This paper |
| JJ341 | *MaHCKRS2* | This paper |
| JJ342 | *MaHCKRS3* | This paper |
| JJ82 | *SL2::HuSmad-4_NLS::PCKRS::let-858_3’UTR* | This paper |
| JJ426 | *mKate2::let-858_3’UTR* | This paper |
| JJ472 | *SL2::4x(rpr-1p::tRNA(M15)::sup-7_3’UTR)::let-858_3’UTR* | This paper |
| JJ540 | *FLP(C189TAG)::HA::EGL-13_NLS* | ^4^ |
| JJ547 | *MaPCC2RS1* | This paper |
| JJ548 | *MaPCC2RS2* | This paper |
| JJ549 | *MaPCC2RS3* | This paper |
| JJ680 | *SL2::4x(rpr-1p::tRNA(Ma10)[UCA]::sup-7_3’UTR)::let-858_3’UTR* | This paper |
| JJ683 | *FLP(C189TGA)::HA::EGL-13_NLS* | This paper |
| KB124 | *HuSmad-4_NLS::PCC2RS* | ^4^ |
| KB126 | *HuSmad-4_NLS::NPYRS* | ^4^ |
| KB182 | *MaPylRS* | This paper |
| KB257 | *FLP(K223TAG)::HA::EGL-13_NLS* | ^4^ |
| KB321 | *FLP(Y343TAG)::HA::EGL-13_NLS* | ^4^ |
| KB442 | *Citrine2::HA::let-858_3’UTR* | ^4^ |
| LD222 | *FRT::lacZ::FRT* | ^3^ |
| SE72 | *sur-5p* | ^3^ |
| SE77 | *maco-1p* | ^3^ |
| SE80 | *sur-5p::HygR::unc-54_3’UTR* | ^4^ |
| SE154 | *HuSmad-4_NLS::MmPCKRS* | ^3^ |
| SG365 | *loxP::lacZ:loxP* | ^3^ |
| SG606 | *let-858_3’UTR* | ^3^ |

**Supplementary Note 1 Protein sequence of *Ma*PylRS variants.**

Highlighted residues represent:
Variant-specific mutations
H227I, Y228P
V42I, K181E, A190V

***Ma*PCKRS1**
MTVKYTDAQIQRLREYGNGTYEQKVFEDLASRDAAFSKEMSVASTDNEKKIKGMIANPSRHGLTQLMNDIADALVAEGFIEVRTPIFISKDALAR**F**TITEDKPLFKQVFWIDEKRALRPML**S**PNL**C**SVMRDLRDHTDGPVKIFEMGSCFRKESHSGMHLEEFTMLNLVDMGPRGDATEVLKNYISVVMKAAGLPDYDLVQEESDVYKETIDVEINGQEVCSAAVGPHYLDAAHDVHEPWSGAGFGLERLLTIREKYSTVKKGGASISYLNGAKIN*

***Ma*PCKRS2**
MTVKYTDAQIQRLREYGNGTYEQKVFEDLASRDAAFSKEMSVASTDNEKKIKGMIANPSRHGLTQLMNDIADALVAEGFIEVRTPIFISKDALAR**F**TITEDKPLFKQVFWIDEKRALRPML**S**PNL**C**SVMRDLRDHTDGPVKIFEMGSCFRKESHSGMHLEEFTMLNLVDMGPRGDATEVLKNYISVVMKAAGLPDYDLVQEESDVYKETIDVEINGQEVCSAAVGP**IP**LDAAHDVHEPWSGAGFGLERLLTIREKYSTVKKGGASISYLNGAKIN*

***Ma*PCKRS3**
MTVKYTDAQIQRLREYGNGTYEQKVFEDLASRDAAFSKEMS**I**ASTDNEKKIKGMIANPSRHGLTQLMNDIADALVAEGFIEVRTPIFISKDALAR**F**TITEDKPLFKQVFWIDEKRALRPML**S**PNL**C**SVMRDLRDHTDGPVKIFEMGSCFRKESHSGMHLEEFTMLNLVDMGPRGDATEVL**E**NYISVVMK**V**AGLPDYDLVQEESDVYKETIDVEINGQEVCSAAVGP**IP**LDAAHDVHEPWSGAGFGLERLLTIREKYSTVKKGGASISYLNGAKIN*

***Ma*HCKRS1**
MTVKYTDAQIQRLREYGNGTYEQKVFEDLASRDAAFSKEMSVASTDNEKKIKGMIANPSRHGLTQLMNDIADALVAEGFIEVRTPIFISKDALARMTITEDKPLFKQVFWIDEKRALRPMLAPNL**A**SVMRDLRDHTDGPVKIFEMGSCFRKESHSGMHLEEFTMLNLVDMGPRGDATEVLKNYISVVMKAAGLPDYDLVQEESDVYKETIDVEINGQEVCSAAVGPHYLDAAHDVHEPWSGAGFGLERLLTIREKYSTVKKGGASISYLNGAKIN*

***Ma*HCKRS2**
MTVKYTDAQIQRLREYGNGTYEQKVFEDLASRDAAFSKEMSVASTDNEKKIKGMIANPSRHGLTQLMNDIADALVAEGFIEVRTPIFISKDALARMTITEDKPLFKQVFWIDEKRALRPMLAPNL**A**SVMRDLRDHTDGPVKIFEMGSCFRKESHSGMHLEEFTMLNLVDMGPRGDATEVLKNYISVVMKAAGLPDYDLVQEESDVYKETIDVEINGQEVCSAAVGP**IP**LDAAHDVHEPWSGAGFGLERLLTIREKYSTVKKGGASISYLNGAKIN*

***Ma*HCKRS3**
MTVKYTDAQIQRLREYGNGTYEQKVFEDLASRDAAFSKEMS**I**ASTDNEKKIKGMIANPSRHGLTQLMNDIADALVAEGFIEVRTPIFISKDALARMTITEDKPLFKQVFWIDEKRALRPMLAPNL**A**SVMRDLRDHTDGPVKIFEMGSCFRKESHSGMHLEEFTMLNLVDMGPRGDATEVL**E**NYISVVMK**V**AGLPDYDLVQEESDVYKETIDVEINGQEVCSAAVGP**IP**LDAAHDVHEPWSGAGFGLERLLTIREKYSTVKKGGASISYLNGAKIN*

***Ma*NPYRS1**
MTVKYTDAQIQRLREYGNGTYEQKVFEDLASRDAAFSKEMSVASTDNEKKIKGMIANPSRHGLTQLMNDIADALVAEGFIEVRTPIFISKDALARMTITEDKPLFKQVFWIDEKRALRPMLAPN**F**YSVMRDLRDHTDGPVKIFEMGSCFRKESHSGMHLEEFTML**G**L**G**DMGPRGDATEVLKNYISVVMKAAGLPDYDLVQEESDV**F**KETIDVEINGQEVCSAAVGPHYLDAAHDVHEPWSGAGFGLERLLTIREKYSTVKKGGASISYLNGAKIN*

***Ma*NPYRS2**
MTVKYTDAQIQRLREYGNGTYEQKVFEDLASRDAAFSKEMSVASTDNEKKIKGMIANPSRHGLTQLMNDIADALVAEGFIEVRTPIFISKDALARMTITEDKPLFKQVFWIDEKRALRPMLAPN**F**YSVMRDLRDHTDGPVKIFEMGSCFRKESHSGMHLEEFTML**G**L**G**DMGPRGDATEVLKNYISVVMKAAGLPDYDLVQEESDV**F**KETIDVEINGQEVCSAAVGP**IP**LDAAHDVHEPWSGAGFGLERLLTIREKYSTVKKGGASISYLNGAKIN*

***Ma*NPYRS3**
MTVKYTDAQIQRLREYGNGTYEQKVFEDLASRDAAFSKEMS**I**ASTDNEKKIKGMIANPSRHGLTQLMNDIADALVAEGFIEVRTPIFISKDALARMTITEDKPLFKQVFWIDEKRALRPMLAPN**F**YSVMRDLRDHTDGPVKIFEMGSCFRKESHSGMHLEEFTML**G**L**G**DMGPRGDATEVL**E**NYISVVMK**V**AGLPDYDLVQEESDV**F**KETIDVEINGQEVCSAAVGP**IP**LDAAHDVHEPWSGAGFGLERLLTIREKYSTVKKGGASISYLNGAKIN*

***Ma*PCC2RS1**
MTVKYTDAQIQRLREYGNGTYEQKVFEDLASRDAAFSKEMSVASTDNEKKIKGMIANPSRHGLTQLMNDIADALVAEGFIEVRTPIFISKDALARMTITEDKPLFKQVFWIDEKRALRPMLAPNLYSVMRDLRDHTDGPVKIFEMGSCFRKESHSGMHLEEFTML**Q**L**A**DMGPRGDATEVLKNYISVVMKAAGLPDYDLVQEESDVYKETIDVEINGQEVCSA**M**VGPHYLDAAHDVHEPWSGAGFGLERLLTIREKYSTVKKGGASISYLNGAKIN*

***Ma*PCC2RS2**
MTVKYTDAQIQRLREYGNGTYEQKVFEDLASRDAAFSKEMSVASTDNEKKIKGMIANPSRHGLTQLMNDIADALVAEGFIEVRTPIFISKDALARMTITEDKPLFKQVFWIDEKRALRPMLAPNLYSVMRDLRDHTDGPVKIFEMGSCFRKESHSGMHLEEFTML**Q**L**A**DMGPRGDATEVLKNYISVVMKAAGLPDYDLVQEESDVYKETIDVEINGQEVCSA**M**VGP**IP**LDAAHDVHEPWSGAGFGLERLLTIREKYSTVKKGGASISYLNGAKIN*

***Ma*PCC2RS3**
MTVKYTDAQIQRLREYGNGTYEQKVFEDLASRDAAFSKEMS**I**ASTDNEKKIKGMIANPSRHGLTQLMNDIADALVAEGFIEVRTPIFISKDALARMTITEDKPLFKQVFWIDEKRALRPMLAPNLYSVMRDLRDHTDGPVKIFEMGSCFRKESHSGMHLEEFTML**Q**L**A**DMGPRGDATEVL**E**NYISVVMK**V**AGLPDYDLVQEESDVYKETIDVEINGQEVCSA**M**VGP**IP**LDAAHDVHEPWSGAGFGLERLLTIREKYSTVKKGGASISYLNGAKIN*

**Supplementary Note 2 *C. elegans* optimised nucleotide sequence of *Ma*PylRS variants.**

Highlighted bases represent:
Synthetic introns

***Ma*PCKRS1**
ATGACAGTAAAGTACACCGATGCCCAGATACAGAGACTCCGTGAGTACGGAAACGGAACCTACGAGCAAAAGGTCTTCGAGGACCTCGCCTCCCGTGACGCCGCCTTCTCCAAGGAGATGTCCGTCGCCTCCACCGACAACGAGAAGAAGATCAAGgtaagtttatacatatatatactaactaaccctgattatttaaattttcagGGAATGATCGCCAACCCATCCCGTCACGGACTCACCCAACTCATGAACGACATCGCCGACGCCCTCGTCGCCGAGGGATTCATCGAGGTCCGTACCCCAATCTTCATCTCCAAGGACGCCCTCGCCCGTtTcACCATCACCGAGGACAAGCCACTCTTCAAGCAAGTCTTCTGGATCGACGAGAAGCGTGCCCTCCGTCCAATGCTCtCCCCAAACCTCTgCTCCGTCATGCGTGACCTCCGTGACCACACCGACGGACCAGTCAAGgtaagtttaatcagttcggtactaactaaccatacatatttaaattttcagATCTTCGAGATGGGATCCTGCTTCCGTAAGGAGTCCCACTCCGGAATGCACCTCGAGGAGTTCACCATGCTCAACCTCGTCGACATGGGACCACGTGGAGACGCCACCGAGGTCCTCAAGAACTACATCTCCGTCGTCATGAAGGCCGCCGGACTCCCAGACTACGACCTCGTCCAAGAGGAGTCCGACGTCTACAAGgtaagtttaaacatgattttactaactaactaatctgatttaaattttcagGAGACCATCGACGTCGAGATCAACGGACAAGAGGTCTGCTCCGCCGCCGTCGGACCACACTACCTCGACGCCGCCCACGACGTCCACGAGCCATGGTCCGGAGCCGGATTCGGACTCGAGCGTCTCCTCACCATCCGTGAGAAGTACTCCACCGTCAAGAAGGGAGGAGCCTCCATCTCCTACCTCAACGGAGCCAAGATCAACTAA

***Ma*PCKRS2**
ATGACAGTAAAGTACACCGATGCCCAGATACAGAGACTCCGTGAGTACGGAAACGGAACCTACGAGCAAAAGGTCTTCGAGGACCTCGCCTCCCGTGACGCCGCCTTCTCCAAGGAGATGTCCGTCGCCTCCACCGACAACGAGAAGAAGATCAAGgtaagtttatacatatatatactaactaaccctgattatttaaattttcagGGAATGATCGCCAACCCATCCCGTCACGGACTCACCCAACTCATGAACGACATCGCCGACGCCCTCGTCGCCGAGGGATTCATCGAGGTCCGTACCCCAATCTTCATCTCCAAGGACGCCCTCGCCCGTtTcACCATCACCGAGGACAAGCCACTCTTCAAGCAAGTCTTCTGGATCGACGAGAAGCGTGCCCTCCGTCCAATGCTCtCCCCAAACCTCTgCTCCGTCATGCGTGACCTCCGTGACCACACCGACGGACCAGTCAAGgtaagtttaatcagttcggtactaactaaccatacatatttaaattttcagATCTTCGAGATGGGATCCTGCTTCCGTAAGGAGTCCCACTCCGGAATGCACCTCGAGGAGTTCACCATGCTCAACCTCGTCGACATGGGACCACGTGGAGACGCCACCGAGGTCCTCAAGAACTACATCTCCGTCGTCATGAAGGCCGCCGGACTCCCAGACTACGACCTCGTCCAAGAGGAGTCCGACGTCTACAAGgtaagtttaaacatgattttactaactaactaatctgatttaaattttcagGAGACCATCGACGTCGAGATCAACGGACAAGAGGTCTGCTCCGCCGCCGTCGGACCAATCCCACTCGACGCCGCCCACGACGTCCACGAGCCATGGTCCGGAGCCGGATTCGGACTCGAGCGTCTCCTCACCATCCGTGAGAAGTACTCCACCGTCAAGAAGGGAGGAGCCTCCATCTCCTACCTCAACGGAGCCAAGATCAACTAA

***Ma*PCKRS3**
ATGACAGTAAAGTACACCGATGCCCAGATACAGAGACTCCGTGAGTACGGAAACGGAACCTACGAGCAAAAGGTCTTCGAGGACCTCGCCTCCCGTGACGCCGCCTTCTCCAAGGAGATGTCCATTGCCTCCACCGACAACGAGAAGAAGATCAAGgtaagtttatacatatatatactaactaaccctgattatttaaattttcagGGAATGATCGCCAACCCATCCCGTCACGGACTCACCCAACTCATGAACGACATCGCCGACGCCCTCGTCGCCGAGGGATTCATCGAGGTCCGTACCCCAATCTTCATCTCCAAGGACGCCCTCGCCCGTtTcACCATCACCGAGGACAAGCCACTCTTCAAGCAAGTCTTCTGGATCGACGAGAAGCGTGCCCTCCGTCCAATGCTCtCCCCAAACCTCTgCTCCGTCATGCGTGACCTCCGTGACCACACCGACGGACCAGTCAAGgtaagtttaatcagttcggtactaactaaccatacatatttaaattttcagATCTTCGAGATGGGATCCTGCTTCCGTAAGGAGTCCCACTCCGGAATGCACCTCGAGGAGTTCACCATGCTCAACCTCGTCGACATGGGACCACGTGGAGACGCCACCGAGGTCCTCGAAAACTACATCTCCGTCGTCATGAAGGTGGCCGGACTCCCAGACTACGACCTCGTCCAAGAGGAGTCCGACGTCTACAAGgtaagtttaaacatgattttactaactaactaatctgatttaaattttcagGAGACCATCGACGTCGAGATCAACGGACAAGAGGTCTGCTCCGCCGCCGTCGGACCAATCCCACTCGACGCCGCCCACGACGTCCACGAGCCATGGTCCGGAGCCGGATTCGGACTCGAGCGTCTCCTCACCATCCGTGAGAAGTACTCCACCGTCAAGAAGGGAGGAGCCTCCATCTCCTACCTCAACGGAGCCAAGATCAACTAA

***Ma*HCKRS1**
ATGACAGTAAAGTACACCGATGCCCAGATACAGAGACTCCGTGAGTACGGAAACGGAACCTACGAGCAAAAGGTCTTCGAGGACCTCGCCTCCCGTGACGCCGCCTTCTCCAAGGAGATGTCCGTCGCCTCCACCGACAACGAGAAGAAGATCAAGgtaagtttatacatatatatactaactaaccctgattatttaaattttcagGGAATGATCGCCAACCCATCCCGTCACGGACTCACCCAACTCATGAACGACATCGCCGACGCCCTCGTCGCCGAGGGATTCATCGAGGTCCGTACCCCAATCTTCATCTCCAAGGACGCCCTCGCCCGTATGACCATCACCGAGGACAAGCCACTCTTCAAGCAAGTCTTCTGGATCGACGAGAAGCGTGCCCTCCGTCCAATGCTCGCCCCAAACCTCGCCTCCGTCATGCGTGACCTCCGTGACCACACCGACGGACCAGTCAAGgtaagtttaatcagttcggtactaactaaccatacatatttaaattttcagATCTTCGAGATGGGATCCTGCTTCCGTAAGGAGTCCCACTCCGGAATGCACCTCGAGGAGTTCACCATGCTCAACCTCGTCGACATGGGACCACGTGGAGACGCCACCGAGGTCCTCAAGAACTACATCTCCGTCGTCATGAAGGCCGCCGGACTCCCAGACTACGACCTCGTCCAAGAGGAGTCCGACGTCTACAAGgtaagtttaaacatgattttactaactaactaatctgatttaaattttcagGAGACCATCGACGTCGAGATCAACGGACAAGAGGTCTGCTCCGCCGCCGTCGGACCACACTACCTCGACGCCGCCCACGACGTCCACGAGCCATGGTCCGGAGCCGGATTCGGACTCGAGCGTCTCCTCACCATCCGTGAGAAGTACTCCACCGTCAAGAAGGGAGGAGCCTCCATCTCCTACCTCAACGGAGCCAAGATCAACTAA

***Ma*HCKRS2**
ATGACAGTAAAGTACACCGATGCCCAGATACAGAGACTCCGTGAGTACGGAAACGGAACCTACGAGCAAAAGGTCTTCGAGGACCTCGCCTCCCGTGACGCCGCCTTCTCCAAGGAGATGTCCGTCGCCTCCACCGACAACGAGAAGAAGATCAAGgtaagtttatacatatatatactaactaaccctgattatttaaattttcagGGAATGATCGCCAACCCATCCCGTCACGGACTCACCCAACTCATGAACGACATCGCCGACGCCCTCGTCGCCGAGGGATTCATCGAGGTCCGTACCCCAATCTTCATCTCCAAGGACGCCCTCGCCCGTATGACCATCACCGAGGACAAGCCACTCTTCAAGCAAGTCTTCTGGATCGACGAGAAGCGTGCCCTCCGTCCAATGCTCGCCCCAAACCTCGCCTCCGTCATGCGTGACCTCCGTGACCACACCGACGGACCAGTCAAGgtaagtttaatcagttcggtactaactaaccatacatatttaaattttcagATCTTCGAGATGGGATCCTGCTTCCGTAAGGAGTCCCACTCCGGAATGCACCTCGAGGAGTTCACCATGCTCAACCTCGTCGACATGGGACCACGTGGAGACGCCACCGAGGTCCTCAAGAACTACATCTCCGTCGTCATGAAGGCCGCCGGACTCCCAGACTACGACCTCGTCCAAGAGGAGTCCGACGTCTACAAGgtaagtttaaacatgattttactaactaactaatctgatttaaattttcagGAGACCATCGACGTCGAGATCAACGGACAAGAGGTCTGCTCCGCCGCCGTCGGACCAATCCCACTCGACGCCGCCCACGACGTCCACGAGCCATGGTCCGGAGCCGGATTCGGACTCGAGCGTCTCCTCACCATCCGTGAGAAGTACTCCACCGTCAAGAAGGGAGGAGCCTCCATCTCCTACCTCAACGGAGCCAAGATCAACTAA

***Ma*HCKRS3**
ATGACAGTAAAGTACACCGATGCCCAGATACAGAGACTCCGTGAGTACGGAAACGGAACCTACGAGCAAAAGGTCTTCGAGGACCTCGCCTCCCGTGACGCCGCCTTCTCCAAGGAGATGTCCATTGCCTCCACCGACAACGAGAAGAAGATCAAGgtaagtttatacatatatatactaactaaccctgattatttaaattttcagGGAATGATCGCCAACCCATCCCGTCACGGACTCACCCAACTCATGAACGACATCGCCGACGCCCTCGTCGCCGAGGGATTCATCGAGGTCCGTACCCCAATCTTCATCTCCAAGGACGCCCTCGCCCGTATGACCATCACCGAGGACAAGCCACTCTTCAAGCAAGTCTTCTGGATCGACGAGAAGCGTGCCCTCCGTCCAATGCTCGCCCCAAACCTCGCCTCCGTCATGCGTGACCTCCGTGACCACACCGACGGACCAGTCAAGgtaagtttaatcagttcggtactaactaaccatacatatttaaattttcagATCTTCGAGATGGGATCCTGCTTCCGTAAGGAGTCCCACTCCGGAATGCACCTCGAGGAGTTCACCATGCTCAACCTCGTCGACATGGGACCACGTGGAGACGCCACCGAGGTCCTCGAAAACTACATCTCCGTCGTCATGAAGGTGGCCGGACTCCCAGACTACGACCTCGTCCAAGAGGAGTCCGACGTCTACAAGgtaagtttaaacatgattttactaactaactaatctgatttaaattttcagGAGACCATCGACGTCGAGATCAACGGACAAGAGGTCTGCTCCGCCGCCGTCGGACCAATCCCACTCGACGCCGCCCACGACGTCCACGAGCCATGGTCCGGAGCCGGATTCGGACTCGAGCGTCTCCTCACCATCCGTGAGAAGTACTCCACCGTCAAGAAGGGAGGAGCCTCCATCTCCTACCTCAACGGAGCCAAGATCAACTAA

***Ma*NPYRS1**
ATGACAGTAAAGTACACCGATGCCCAGATACAGAGACTCCGTGAGTACGGAAACGGAACCTACGAGCAAAAGGTCTTCGAGGACCTCGCCTCCCGTGACGCCGCCTTCTCCAAGGAGATGTCCGTCGCCTCCACCGACAACGAGAAGAAGATCAAGgtaagtttatacatatatatactaactaaccctgattatttaaattttcagGGAATGATCGCCAACCCATCCCGTCACGGACTCACCCAACTCATGAACGACATCGCCGACGCCCTCGTCGCCGAGGGATTCATCGAGGTCCGTACCCCAATCTTCATCTCCAAGGACGCCCTCGCCCGTATGACCATCACCGAGGACAAGCCACTCTTCAAGCAAGTCTTCTGGATCGACGAGAAGCGTGCCCTCCGTCCAATGCTCGCCCCAAACttcTACTCCGTCATGCGTGACCTCCGTGACCACACCGACGGACCAGTCAAGgtaagtttaatcagttcggtactaactaaccatacatatttaaattttcagATCTTCGAGATGGGATCCTGCTTCCGTAAGGAGTCCCACTCCGGAATGCACCTCGAGGAGTTCACCATGCTCggaCTCggaGACATGGGACCACGTGGAGACGCCACCGAGGTCCTCAAGAACTACATCTCCGTCGTCATGAAGGCCGCCGGACTCCCAGACTACGACCTCGTCCAAGAGGAGTCCGACGTCTTCAAGgtaagtttaaacatgattttactaactaactaatctgatttaaattttcagGAGACCATCGACGTCGAGATCAACGGACAAGAGGTCTGCTCCGCCGCCGTCGGACCACACTACCTCGACGCCGCCCACGACGTCCACGAGCCATGGTCCGGAGCCGGATTCGGACTCGAGCGTCTCCTCACCATCCGTGAGAAGTACTCCACCGTCAAGAAGGGAGGAGCCTCCATCTCCTACCTCAACGGAGCCAAGATCAACTAA

***Ma*NPYRS2**
ATGACAGTAAAGTACACCGATGCCCAGATACAGAGACTCCGTGAGTACGGAAACGGAACCTACGAGCAAAAGGTCTTCGAGGACCTCGCCTCCCGTGACGCCGCCTTCTCCAAGGAGATGTCCGTCGCCTCCACCGACAACGAGAAGAAGATCAAGgtaagtttatacatatatatactaactaaccctgattatttaaattttcagGGAATGATCGCCAACCCATCCCGTCACGGACTCACCCAACTCATGAACGACATCGCCGACGCCCTCGTCGCCGAGGGATTCATCGAGGTCCGTACCCCAATCTTCATCTCCAAGGACGCCCTCGCCCGTATGACCATCACCGAGGACAAGCCACTCTTCAAGCAAGTCTTCTGGATCGACGAGAAGCGTGCCCTCCGTCCAATGCTCGCCCCAAACttcTACTCCGTCATGCGTGACCTCCGTGACCACACCGACGGACCAGTCAAGgtaagtttaatcagttcggtactaactaaccatacatatttaaattttcagATCTTCGAGATGGGATCCTGCTTCCGTAAGGAGTCCCACTCCGGAATGCACCTCGAGGAGTTCACCATGCTCggaCTCggaGACATGGGACCACGTGGAGACGCCACCGAGGTCCTCAAGAACTACATCTCCGTCGTCATGAAGGCCGCCGGACTCCCAGACTACGACCTCGTCCAAGAGGAGTCCGACGTCTTCAAGgtaagtttaaacatgattttactaactaactaatctgatttaaattttcagGAGACCATCGACGTCGAGATCAACGGACAAGAGGTCTGCTCCGCCGCCGTCGGACCAATCCCACTCGACGCCGCCCACGACGTCCACGAGCCATGGTCCGGAGCCGGATTCGGACTCGAGCGTCTCCTCACCATCCGTGAGAAGTACTCCACCGTCAAGAAGGGAGGAGCCTCCATCTCCTACCTCAACGGAGCCAAGATCAACTAA

***Ma*NPYRS3**
ATGACAGTAAAGTACACCGATGCCCAGATACAGAGACTCCGTGAGTACGGAAACGGAACCTACGAGCAAAAGGTCTTCGAGGACCTCGCCTCCCGTGACGCCGCCTTCTCCAAGGAGATGTCCATTGCCTCCACCGACAACGAGAAGAAGATCAAGgtaagtttatacatatatatactaactaaccctgattatttaaattttcagGGAATGATCGCCAACCCATCCCGTCACGGACTCACCCAACTCATGAACGACATCGCCGACGCCCTCGTCGCCGAGGGATTCATCGAGGTCCGTACCCCAATCTTCATCTCCAAGGACGCCCTCGCCCGTATGACCATCACCGAGGACAAGCCACTCTTCAAGCAAGTCTTCTGGATCGACGAGAAGCGTGCCCTCCGTCCAATGCTCGCCCCAAACttcTACTCCGTCATGCGTGACCTCCGTGACCACACCGACGGACCAGTCAAGgtaagtttaatcagttcggtactaactaaccatacatatttaaattttcagATCTTCGAGATGGGATCCTGCTTCCGTAAGGAGTCCCACTCCGGAATGCACCTCGAGGAGTTCACCATGCTCggaCTCggaGACATGGGACCACGTGGAGACGCCACCGAGGTCCTCGAAAACTACATCTCCGTCGTCATGAAGGTGGCCGGACTCCCAGACTACGACCTCGTCCAAGAGGAGTCCGACGTCTTCAAGgtaagtttaaacatgattttactaactaactaatctgatttaaattttcagGAGACCATCGACGTCGAGATCAACGGACAAGAGGTCTGCTCCGCCGCCGTCGGACCAATCCCACTCGACGCCGCCCACGACGTCCACGAGCCATGGTCCGGAGCCGGATTCGGACTCGAGCGTCTCCTCACCATCCGTGAGAAGTACTCCACCGTCAAGAAGGGAGGAGCCTCCATCTCCTACCTCAACGGAGCCAAGATCAACTAA

***Ma*PCC2RS1**
ATGACAGTAAAGTACACCGATGCCCAGATACAGAGACTCCGTGAGTACGGAAACGGAACCTACGAGCAAAAGGTCTTCGAGGACCTCGCCTCCCGTGACGCCGCCTTCTCCAAGGAGATGTCCGTCGCCTCCACCGACAACGAGAAGAAGATCAAGgtaagtttaTacatatatatactaactaaccctgattatttaaattttcagGGAATGATCGCCAACCCATCCCGTCACGGACTCACCCAACTCATGAACGACATCGCCGACGCCCTCGTCGCCGAGGGATTCATCGAGGTCCGTACCCCAATCTTCATCTCCAAGGACGCCCTCGCCCGTATGACCATCACCGAGGACAAGCCACTCTTCAAGCAAGTCTTCTGGATCGACGAGAAGCGTGCCCTCCGTCCAATGCTCGCCCCAAACCTCTACTCCGTCATGCGTGACCTCCGTGACCACACCGACGGACCAGTCAAGgtaagtttaaTcagttcggtactaactaaccatacatatttaaattttcagATCTTCGAGATGGGATCCTGCTTCCGTAAGGAGTCCCACTCCGGAATGCACCTCGAGGAGTTCACCATGCTCCAACTCGCCGACATGGGACCACGTGGAGACGCCACCGAGGTCCTCAAGAACTACATCTCCGTCGTCATGAAGGCCGCCGGACTCCCAGACTACGACCTCGTCCAAGAGGAGTCCGACGTCTACAAGgtaagtttaaacatgattttactaactaactaatctgatttaaattttcagGAGACCATCGACGTCGAGATCAACGGACAAGAGGTCTGCTCCGCCATGGTCGGACCACACTACCTCGACGCCGCCCACGACGTCCACGAGCCATGGTCCGGAGCCGGATTCGGACTCGAGCGTCTCCTCACCATCCGTGAGAAGTACTCCACCGTCAAGAAGGGAGGAGCCTCCATCTCCTACCTCAACGGAGCCAAGATCAACTAA

***Ma*PCC2RS2**
ATGACAGTAAAGTACACCGATGCCCAGATACAGAGACTCCGTGAGTACGGAAACGGAACCTACGAGCAAAAGGTCTTCGAGGACCTCGCCTCCCGTGACGCCGCCTTCTCCAAGGAGATGTCCGTCGCCTCCACCGACAACGAGAAGAAGATCAAGgtaagtttaTacatatatatactaactaaccctgattatttaaattttcagGGAATGATCGCCAACCCATCCCGTCACGGACTCACCCAACTCATGAACGACATCGCCGACGCCCTCGTCGCCGAGGGATTCATCGAGGTCCGTACCCCAATCTTCATCTCCAAGGACGCCCTCGCCCGTATGACCATCACCGAGGACAAGCCACTCTTCAAGCAAGTCTTCTGGATCGACGAGAAGCGTGCCCTCCGTCCAATGCTCGCCCCAAACCTCTACTCCGTCATGCGTGACCTCCGTGACCACACCGACGGACCAGTCAAGgtaagtttaaTcagttcggtactaactaaccatacatatttaaattttcagATCTTCGAGATGGGATCCTGCTTCCGTAAGGAGTCCCACTCCGGAATGCACCTCGAGGAGTTCACCATGCTCCAACTCGCCGACATGGGACCACGTGGAGACGCCACCGAGGTCCTCAAGAACTACATCTCCGTCGTCATGAAGGCCGCCGGACTCCCAGACTACGACCTCGTCCAAGAGGAGTCCGACGTCTACAAGgtaagtttaaacatgattttactaactaactaatctgatttaaattttcagGAGACCATCGACGTCGAGATCAACGGACAAGAGGTCTGCTCCGCCATGGTCGGACCA

***Ma*PCC2RS3**
ATGACAGTAAAGTACACCGATGCCCAGATACAGAGACTCCGTGAGTACGGAAACGGAACCTACGAGCAAAAGGTCTTCGAGGACCTCGCCTCCCGTGACGCCGCCTTCTCCAAGGAGATGTCCATTGCCTCCACCGACAACGAGAAGAAGATCAAGgtaagtttaTacatatatatactaactaaccctgattatttaaattttcagGGAATGATCGCCAACCCATCCCGTCACGGACTCACCCAACTCATGAACGACATCGCCGACGCCCTCGTCGCCGAGGGATTCATCGAGGTCCGTACCCCAATCTTCATCTCCAAGGACGCCCTCGCCCGTATGACCATCACCGAGGACAAGCCACTCTTCAAGCAAGTCTTCTGGATCGACGAGAAGCGTGCCCTCCGTCCAATGCTCGCCCCAAACCTCTACTCCGTCATGCGTGACCTCCGTGACCACACCGACGGACCAGTCAAGgtaagtttaaTcagttcggtactaactaaccatacatatttaaattttcagATCTTCGAGATGGGATCCTGCTTCCGTAAGGAGTCCCACTCCGGAATGCACCTCGAGGAGTTCACCATGCTCCAACTCGCCGACATGGGACCACGTGGAGACGCCACCGAGGTCCTCGAAAACTACATCTCCGTCGTCATGAAGGTGGCCGGACTCCCAGACTACGACCTCGTCCAAGAGGAGTCCGACGTCTACAAGgtaagtttaaacatgattttactaactaactaatctgatttaaattttcagGAGACCATCGACGTCGAGATCAACGGACAAGAGGTCTGCTCCGCCATGGTCGGACCAATCCCACTCGACGCCGCCCACGACGTCCACGAGCCATGGTCCGGAGCCGGATTCGGACTCGAGCGTCTCCTCACCATCCGTGAGAAGTACTCCACCGTCAAGAAGGGAGGAGCCTCCATCTCCTACCTCAACGGAGCCAAGATCAACTAA
